# AIRBP: Accurate identification of RNA-binding proteins using machine learning techniques

**DOI:** 10.1101/2020.03.10.985416

**Authors:** Avdesh Mishra, Reecha Khanal, Md Tamjidul Hoque

## Abstract

**Motivation:** Identification of RNA-binding proteins (RBPs) that bind to ribonucleic acid molecules, is an important problem in Computational Biology and Bioinformatics. It becomes indispensable to identify RBPs as they play crucial roles in post-transcriptional control of RNAs and RNA metabolism as well as have diverse roles in various biological processes such as splicing, mRNA stabilization, mRNA localization, and translation, RNA synthesis, folding-unfolding, modification, processing, and degradation. The existing experimental techniques for identifying RBPs are time-consuming and expensive. Therefore, identifying RBPs directly from the sequence using computational methods can be useful to efficiently annotate RBPs and assist the experimental design. In this work, we present a method, called AIRBP, which is designed using an advanced machine learning technique, called stacking, to effectively predict RBPs by utilizing features extracted from evolutionary information, physiochemical properties, and disordered properties. Moreover, our method, AIRBP is trained on the useful feature-subset identified by the evolutionary algorithm (EA).

**Results:** The results show that AIRBP attains Accuracy (ACC), F1-score, and MCC of 95.38%, 0.917, and 0.885, respectively, based on the benchmark dataset, using 10-fold cross-validation (CV). Further evaluation of AIRBP on independent test set reveals that it achieves ACC, F1-score, and MCC of 93.04%, 0.943, and 0.855, for Human test set; 91.60%, 0.942 and 0.789 for S. cerevisiae test set; and 91.67%, 0.953 and 0.594 for A. thaliana test set, respectively. These results indicate that AIRBP outperforms the current state-of-the-art method. Therefore, the proposed top-performing AIRBP can be useful for accurate identification and annotation of RBPs directly from the sequence and help gain valuable insight to treat critical diseases.

**Availability:** Code-data is available here: http://cs.uno.edu/~tamjid/Software/AIRBP/code_data.zip

## Introduction

RNA Binding Proteins (RBPs) are proteins that bind to ribonucleic acid (RNA) molecules and form dynamic units, called ribonucleoprotein (RNP) complexes. These RBPs along with the RNP complexes, play a crucial role starting from the biogenesis process of RNA to its degradation (Beckmann, et al., 2015). Additionally, they contribute to several essential biological functions that include RNA transport, cellular localization, gene expression, expression of histone genes, post-transcriptional gene regulation, and regulation of translation and transcription control (Glisovic, et al., 2008). As an illustration, the newly formed messenger RNA, that carries necessary genetic information from DNA to ribosomes, associates with various RNA binding proteins (RBP) to form messenger ribonucleoprotein (mRNP) complexes (Baltz, et al., 2012). These mRNP complexes govern major elements of the metabolism and functions of mRNA. Similarly, the microRNPs (miRNPs), formed through association of the RBPs with microRNAs (miRNAs) controls the translation and stability of RNA itself (Wurth, 2012). The identification of RBPs along with their mRNA targets, is shown useful in cancer therapy (Wurth, 2012). Numerous other diseases have been linked to defective RBP expression and functions. Some of those diseases are neuropathies, muscular atrophies, and metabolic disorders (Castello, et al., 2012). All this information highlight the urgency of identifying the possible RBPs.

As of today, numerous studies have been performed, and various experimental and computational methods have been developed to identify and expand our knowledge of RBP. The initial steps towards identification and study of RBPs and RNP complexes date back to almost half a century ago where experimental methods such as purification of mRNPs from in vitro UV-irradiated polysomal fractions (Greenberg, 1979), from UV-irradiated intact cells (Wagenmakers, et al., 1980) and untreated cells (Lindberg and Sundquist, 1974) revealed the association of a specific set of proteins with mRNA (Baltz, et al., 2012). Recently, cutting-edge experimental approaches are developed to recognize numerous RBPs, which include identification of 860 RBPs in human HeLa cells (Castello, et al., 2012) using UV crosslinking methods, 797 RBPs in human embryonic kidney cell line (Baltz, et al., 2012) using photoreactive nucleotide-enhanced UV crosslinking and oligo(dT) purification approach, 555 mRNA-binding proteins from mouse embryonic stem cells (Kwon, et al., 2013) using UV crosslinking, oligo(dT) and Mass Spectrometry and 120 RBPs from S. *cerevisiae* cells (Mitchell, et al., 2013) using UV crosslinking and purification methods. These experiments for identifying and analyzing of RBPs, have broadened our understanding of RBPs to a certain extent. Despite the great efforts and achievements, these experiments are expensive, time-consuming and labor-intensive (Si, et al., 2015). Moreover, the tremendous progress in genome sequencing has resulted in an unprecedented amount of genetic information and provided a plethora of protein sequences (Wu, et al., 2006), which outpace the tasks of annotating them and elucidating their functions. Thus, it becomes urgent to have faster and more accurate computational approaches to build an RBP repository and RNA-RBP interaction network maps.

In the recent past, several attempts have been made in identifying RNA-binding proteins and many effective computational prediction methods have been developed, which can be divided into two broad categories: *i)* templated based; and *ii)* machine learning-based. Template-based methods extract significant structural or sequence similarity between the query and a template known to bind RNA to assess the RNA-binding preference of the target sequence (Yang, et al., 2012; Zhao, et al., 2011; Zhao, et al., 2011). Unlike template-based methods, in machine learning methods, the predictive model is created to predict by finding a pattern in the input feature space (Kumar, et al., 2011; Paz, et al., 2016; Shazman and Mandel-Gutfreund, 2008). The machine learning approaches vary in the features employed and the classification algorithm used.

Zhao *et al*. proposed two template-based approaches for predicting RBPs, of which, SPOT-stru (Zhao, et al., 2011) is a structure-based approach, and SPOT-seq (Zhao, et al., 2011) is a sequence-based approach. In SPOT-stru, the relative structural similarity in the form of Z-score and a statistical energy function DFIRE is used to predict RBPs. The results indicate that SPOT-stru achieved the MCC of 0.57 on the benchmark data of 212 RNA-binding domains and 6761 non-RNA binding domains. On the other hand, in SPOT-seq, the fold recognition between the target sequence and template structures using the defined sequence-structure matching score is used to predict RBPs. As shown, SPOT-seq achieved the MCC of 0.62 on the benchmark data of 215 RBP chains and 5765 non-binding protein chains.

The machine learning-based approach for the prediction of RNA-binding proteins involves two crucial steps: *i)* extraction of relevant features, and *ii)* selection of an appropriate classification algorithm. Furthermore, depending on the feature extraction mechanism, the existing predictive method can be segmented into two different categories: *i)* extraction of relevant features from the structure of protein (Paz, et al., 2016; Shazman and Mandel-Gutfreund, 2008); and *ii)* extraction of relevant features from protein sequence (Kumar, et al., 2011; Ma, et al., 2015; Ma, et al., 2015; Zhang and Liu, 2017). BindUp (Paz, et al., 2016) available as a web server, is one of the recent structure-based methods that extract electrostatic features and other properties from the structure of the protein and uses SVM classifier for RBPs prediction. As reported, BindUp attains sensitivity of 0.71 and specificity of 0.96 on an independent test set of 323 structures of RNA binding proteins and a control set of an equal number extracted from Protein Data Bank (PDB). Towards a sequence-based approach, Ma *et al*. (Ma, et al., 2015; Ma, et al., 2015) recently proposed two different methods, which differ in the features used to train the random forest model for predicting. In (Ma, et al., 2015), the authors incorporated features of evolutionary information combined with physicochemical features (EIPP) and amino acid composition feature to develop the random forest predictor. Besides, in (Ma, et al., 2015), the authors employed features such as a conjoint triad, binding propensity, non-binding propensity, and EIPP to establish random forest-based predictor with the minimum redundancy maximum relevance (mRMR) method, followed by incremental feature selection (IFS). As reported, their method achieved an accuracy of 0.8662 and MCC of 0.737. Zhang and Liu (Zhang and Liu, 2017) proposed a new sequence-based approach, namely RBPPred which, integrates the physiochemical properties with the evolutionary information extracted from Position Specific Scoring Matrix (PSSM) profile and utilizes SVM to predict RBPs. As shown, RBPPred correctly predicted 83% of 2780 RBPs and 96% of 7093 non-RBPs with MCC of 0.808 using the 10-fold cross-validation (CV) approach. Despite significant progress, most of the approaches for RBPs prediction developed in the past are limited in explaining how protein-RNA interactions occur. Thus, it is essential to identify new features, effective encoding technique and advanced machine learning techniques that can help further improve the accuracy of RBPs predictor and ultimately improve our understanding of RNA-protein interactions and their functions.

In this work, we explore different sequence-based features, encoding techniques, and machine learning approaches to further improve the prediction accuracy of RNA-binding proteins and our understanding of the binding mechanism of RNA-protein interaction. We propose a method, AIRBP, which utilizes features: Evolutionary Information (EI), Physiochemical Properties (PP), and Disordered Properties (DP). It uses four different types of feature encoding technique: Composition, Transition and Distribution (C-T-D) (Zhang and Liu, 2017), Conjoint Triad (CT) (Wang, et al., 2013; Zhang and Liu, 2017), PSSM Distance Transformation (PSSM-DT) (Mishra, et al., 2018; Xu, et al., 2015) and Residue-wise Contact Energy Matrix Transformation (RCEM-T) (Mishra, et al., 2018). Furthermore, AIRBP utilizes an ensemble machine learning framework, known as stacking (Wolpert, 1992) to predict RBPs from protein sequence only. AIRBP offers a significant improvement in the prediction of RBPs based on the benchmark and independent test datasets when compared to the existing start-of-the-art predictors. Therefore, our predictor can be trusted and used by the research community to guide further the experiments related to RNA-protein interactions and their functions. Further, our study highlights the importance of adding features that account for intrinsically disordered regions in predicting RNA-binding proteins. Our research supports the claim that RNA-binding proteins bind with RNA not only through classical structured RNA-binding domains but also through intrinsically disordered regions. Additionally, our study suggests that the research community would be benefited by considering intrinsically disordered regions in protein that induce binding with RNA, in their experimental studies. We believe that the superior performance of AIRBP will motivate the researchers to use it to identify RNA-binding proteins from sequence information. Moreover, the proposed stacking based machine learning technique, encoding techniques and features discussed in this work could be applied to tackle other relevant biological problems.

## Materials and methods

In this section, we describe the approach for benchmark and independent test data preparation, feature extraction and encoding, performance evaluation metrics, and finally, the path we took to establish the machine learning framework for RBPs prediction.

### Dataset

#### Benchmark dataset

For this work, we collected the updated version of the benchmark dataset first proposed by (<u>Liu</u>; Zhang and Liu, 2017) from web link http://rnabinding.com/RBPPred.html. The updated benchmark dataset was created by the authors (Zhang and Liu, 2017) from the original benchmark dataset by removing 16 proteins that had RNAs in their crystal structure from the negative set. Therefore, the updated benchmark dataset, we collected, consists of 7077 non-RBPs (16 proteins removed from the original benchmark dataset which contained 7093 non-RBPs) and 2780 RBPs (same as the original benchmark dataset). Next, we found that 13 out of 2780 and 90 out of 7077 protein sequences in RBPs and non-RBPs set respectively, contained non-standard amino acids (amino acids other than the 20 standard amino acids). These sequences containing non-standard amino acids were removed from further consideration as the physiochemical properties of non-standard amino acids could not be obtained. Finally, the benchmark dataset which contains 2767 RBPs and 6987 non-RBPs was collected and used for validation and model creation of AIRBP.

#### Independent test set

For this work, we collected the updated version of the benchmark dataset first proposed by (<u>Liu</u>; Zhang and Liu, 2017) from web link http://rnabinding.com/RBPPred.html. This dataset consists of independent test sets for 3 species, human, S*accharomyces cerevisiae* (S. *cerevisiae*) and A*rabidopsis thaliana* (A. *thaliana*). The test set was created by the authors (Zhang and Liu, 2017) from the original independent test set by removing 9 proteins from the human set and 7 proteins from S. *cerevisiae* set that had RNAs in their crystal structure from the negative set, respectively. The updated independent test sets contained a total of 967 RBPs and 588 non-RBPs for human, 354 RBPs and 135 non-RBPs for S. c*erevisiae* and 456 RBPs and 37 non-RBPs for A. *thaliana*. Next, we removed the protein sequences containing non-standard amino acid from each of these independent datasets and finally obtained 967 RBPs and 584 non-RBPs for human, 354 RBPs and 134 non-RBPs for S. *cerevisiae* and 456 RBPs and 36 non-RBPs for A. *thaliana*.

### Feature extraction

To create an effective RBPs predictor from sequence alone, the feature vector for each protein sequence was derived from the PSSM profile, Physiochemical Properties (PP), Residue-wise Contact Energy Matrix (RCEM) and Molecular Recognition Features (MoRFs). A total of 10 different properties was encoded with a vector of 2603 dimensions to represent a protein sequence, as shown in Fig 1. Out of 10, five distinct properties hydrophobicity, polarity, normalized van der Waals volume, polarizability and predicted secondary structure that belongs to PP group were each encoded via 21 dimension vector utilizing the C-T-D encoding technique (Dubchak, et al., 1995; Zhang and Liu, 2017). Moreover, the remaining five properties solvent accessibility, charge and polarity of the side chain, MoRFs, RCEM, and PSSM profile were encoded via 13, 64, 1, 20 and 2400 dimensional vectors, respectively. Here, PSSM belongs to the EI group and MoRFs and RCEM belong to the DP group. The properties solvent accessibility, charge and polarity of the side chain, RCEM and PSSM profile were encoded utilizing C-T-D, CT (Wang, et al., 2013; Zhang and Liu, 2017), RCEM transformation (Mishra, et al., 2018) and PSSM-DT transformation techniques (Mishra, et al., 2018; Xu, et al., 2015), respectively. Each of the 10 properties along with their encoding mechanism is described next in detail.

**Fig 1.**
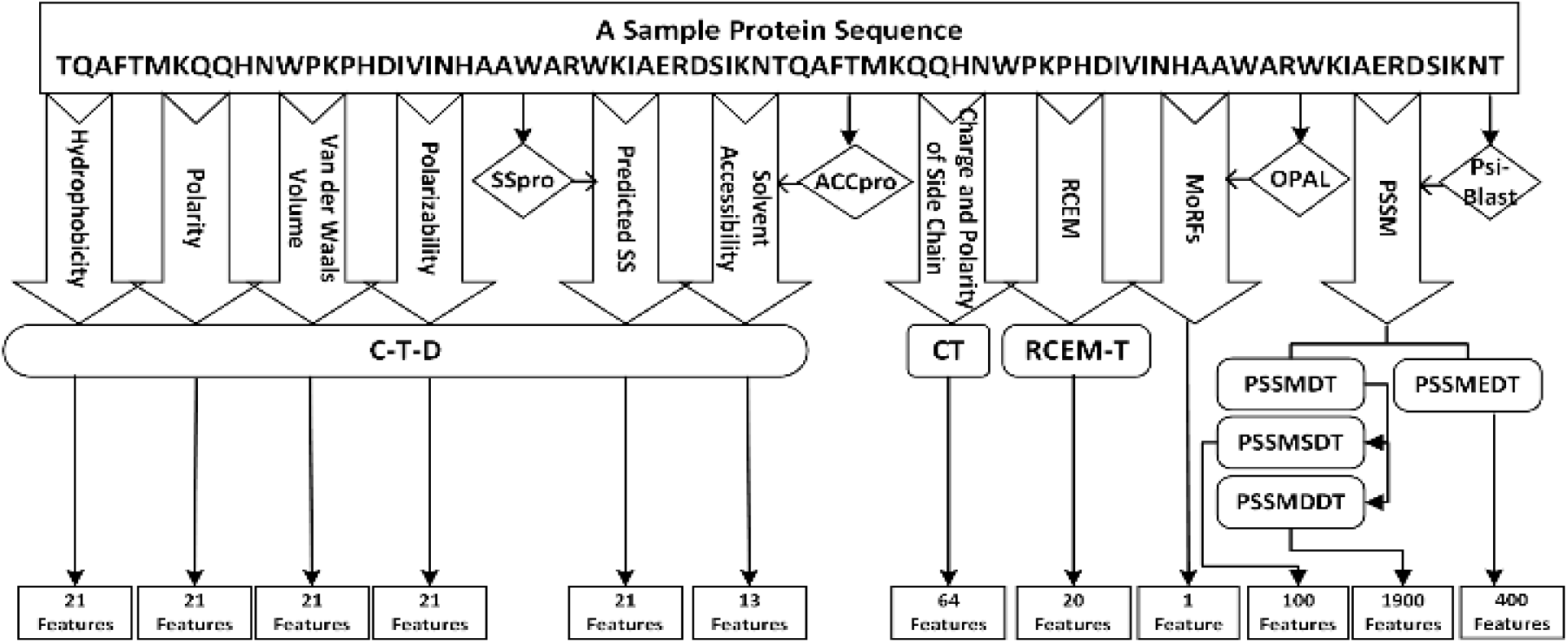
Illustration of encoding the protein sequence into a feature vector of 2603 features utilizing various feature encoding technique. Here, the predicted SS and surface accessibility was obtained from SSpro and ACCpro program (Magnan and Baldi, 2014). Likewise, the MoRFs scores were predicted using the OPAL program (Sharma, et al., 2018) and the PSSM scores were obtained using the PSI-BLAST program (Altschul, et al., 1990)

#### Features extracted from physicochemical properties

In this section we describe various feature extraction techniques, we utilized to obtain a fixed dimensional feature vector from the physicochemical properties which include hydrophobicity, polarity, normalized van der Waals volume, polarizability, predicted secondary structure, solvent accessibility and charge and polarity of the side chain to encode protein sequence.

##### Composition, Transition and Distribution (C-T-D) transformation features

In this section, the C-T-D transformation method aims to describe the distribution patterns of amino acid properties. This method to compute distribution patterns of amino acid properties were first suggested by (Dubchak, et al., 1995) for protein fold class prediction. In our implementation, we used C-T-D transformation to encode the properties including hydrophobicity, polarity, normalized van der Waals volume, polarizability, predicted the secondary structure and solvent accessibility. As the name suggests, this transformation technique focuses on three different components: composition of a particular amino acid in the sequence, transition of one amino acid to other as we go linearly through the sequence, and distribution referring to how one amino acid group is distributed throughout the protein sequence (Han, et al., 2004; Zhang and Liu, 2017). To create a consistent number of features for proteins with different sequence length, 20 standard amino acids are divided into 3 groups (Dubchak, et al., 1999) based on their hydrophobicity, normalized van der Waals volume, polarity, and polarizability. Fig 2 illustrates the C-T-D transformation technique while the 20 standard amino acids are divided into 2 groups which, generates a feature vector of 13 dimensions. Following the transformation, the technique is shown in Fig 2 with an exception that the 20 standard amino acids are divided into 3 groups, we obtain a feature vector of 21 dimensions for the physiochemical properties such as hydrophobicity, normalized van der Waals volume, polarity, and polarizability.

**Fig 2.**
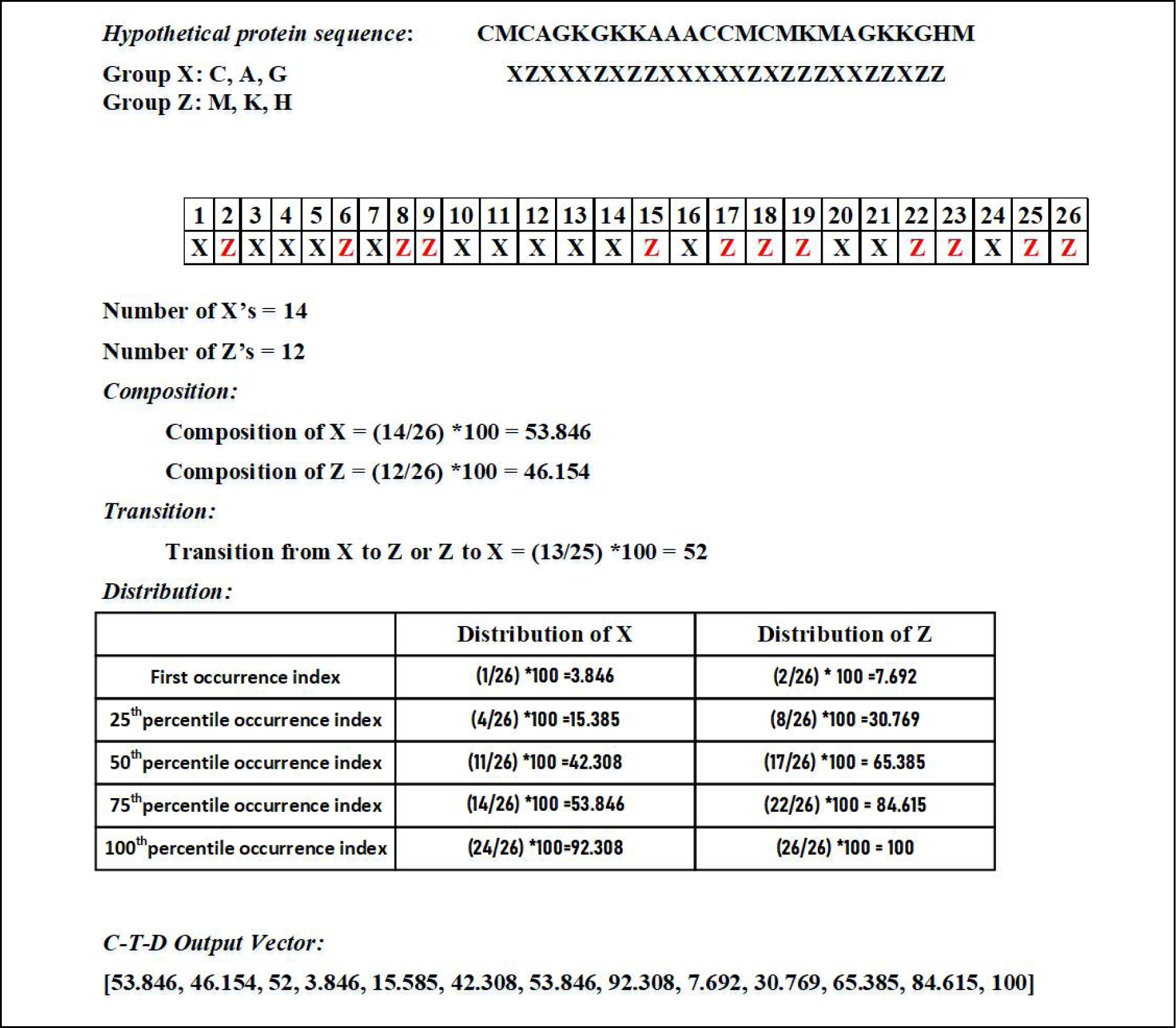
Illustration of the C-T-D transformation technique. While the 20 standard amino acids are divided into 2 groups (e.g., X and Z). First, the group index (X or Z) of every amino acid in the protein sequence is extracted and consequently, a vector of 13 dimensions is obtained through composition, transition, and distribution.

Furthermore, to encode the predicted secondary structure and solvent accessibility as features, we first used the SSpro and ACCpro program (Magnan and Baldi, 2014) to predict secondary structure in the form of ‘H’ (helix), ‘E’ (strand) and ‘C’ (other than helix and strand) and solvent accessibility in the form of ‘e’ (exposed residues) and ‘-’ (buried residues), respectively. The choice of SSpro and ACCpro was made to extract predicted secondary structure and solvent accessibility because of its superior performance and remarkable speed. As reported, SSpro and ACCpro (Magnan and Baldi, 2014) achieved an accuracy of 92.9% and 90% for secondary structure prediction and relative solvent accessibility prediction, respectively. Using the transformation technique described above, we obtained a feature vector of 21 dimensional and 13 dimensions for predicted secondary structure and solvent accessibility, respectively.

##### Conjoint Triad (CT) transformation features

While the 20 standard amino acids are divided into 4 groups (Group A, B, C, and D representing acidic, basic, polar and non-polar, respectively). Shen *et al*. first proposed the CT transformation technique for protein-protein interaction prediction (Shen, et al., 2007), which was successfully applied for protein-RNA interaction prediction in the past (Wang, et al., 2013; Zhang and Liu, 2017). In our implementation, we adopted the CT transformation technique to encode the protein sequence based on the charge and polarity of the side chain of the amino acids in a protein. First, the 20 standard amino acids are divided into 4 groups: *i)* acidic (contain residues D and E); *ii)* basic (contain residues H, R and K); *iii)* polar (contain residues C, G, N, Q, S, T, and Y); and *iv)* non-polar (contain residues A, F, I, L, M, P, V, and W) according to their charge and polarity of the side chain. Then, the protein sequence is converted into a sequence of group types where each element in the sequence represents a group type of the corresponding amino acid in the protein sequence. Next, a triad of three contiguous amino acids is considered as a single unit. Accordingly, all the triads can be classified into 4 × 4 × 4 = 64 classes. Finally, a sliding window of a triad is passed through a sequence of group types and the frequency of occurrences of each type of triad is counted. Through this process, we obtain a feature vector of 64 dimensions for charge and polarity of side chains of amino acids in a protein. Fig 3 provides an illustration of the CT transformation technique we used to extract features from protein sequence based on charge and polarity of side chains.

**Fig 3.**
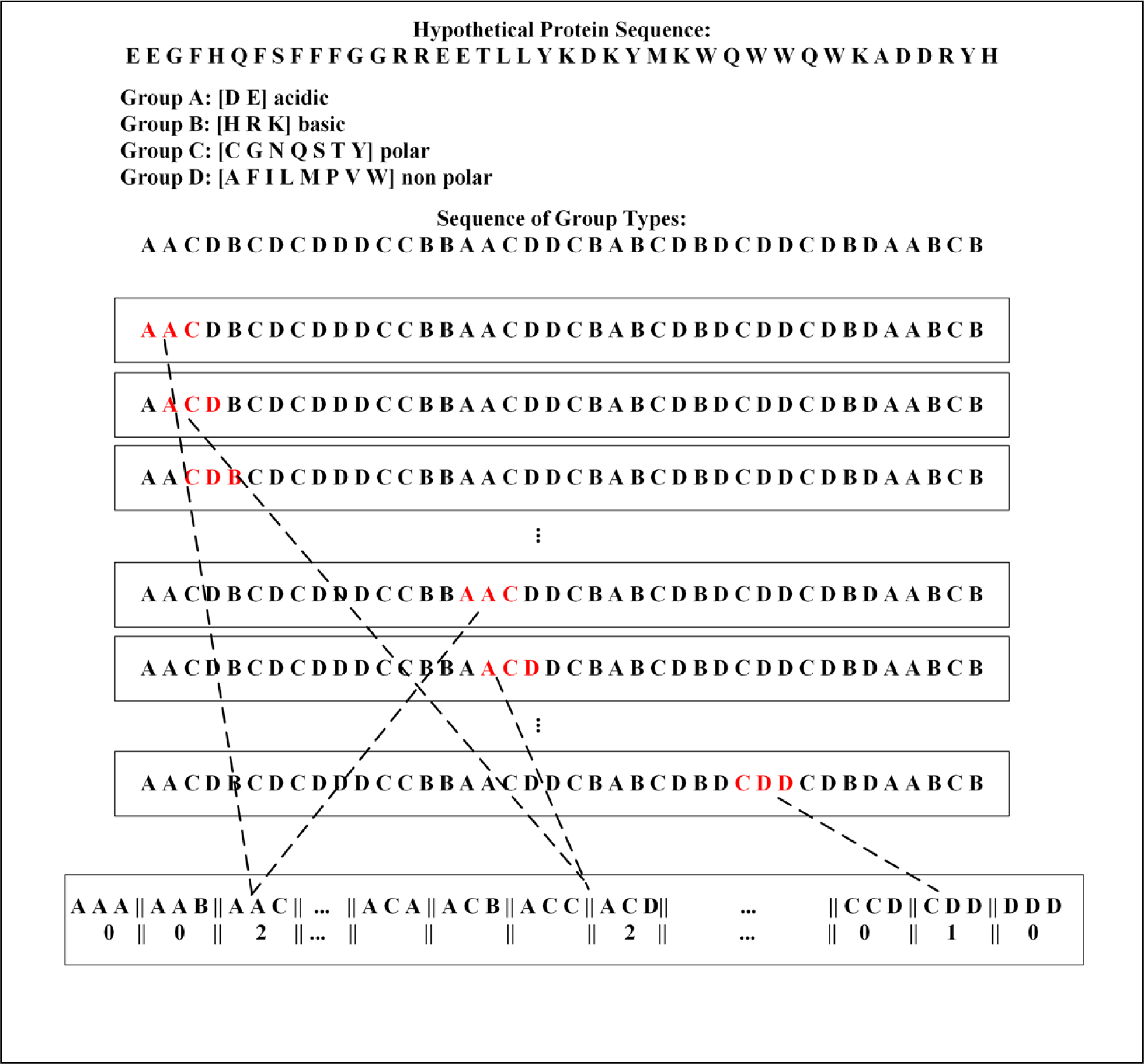
Illustration of Conjoint Triad transformation technique.

**Fig 4.**
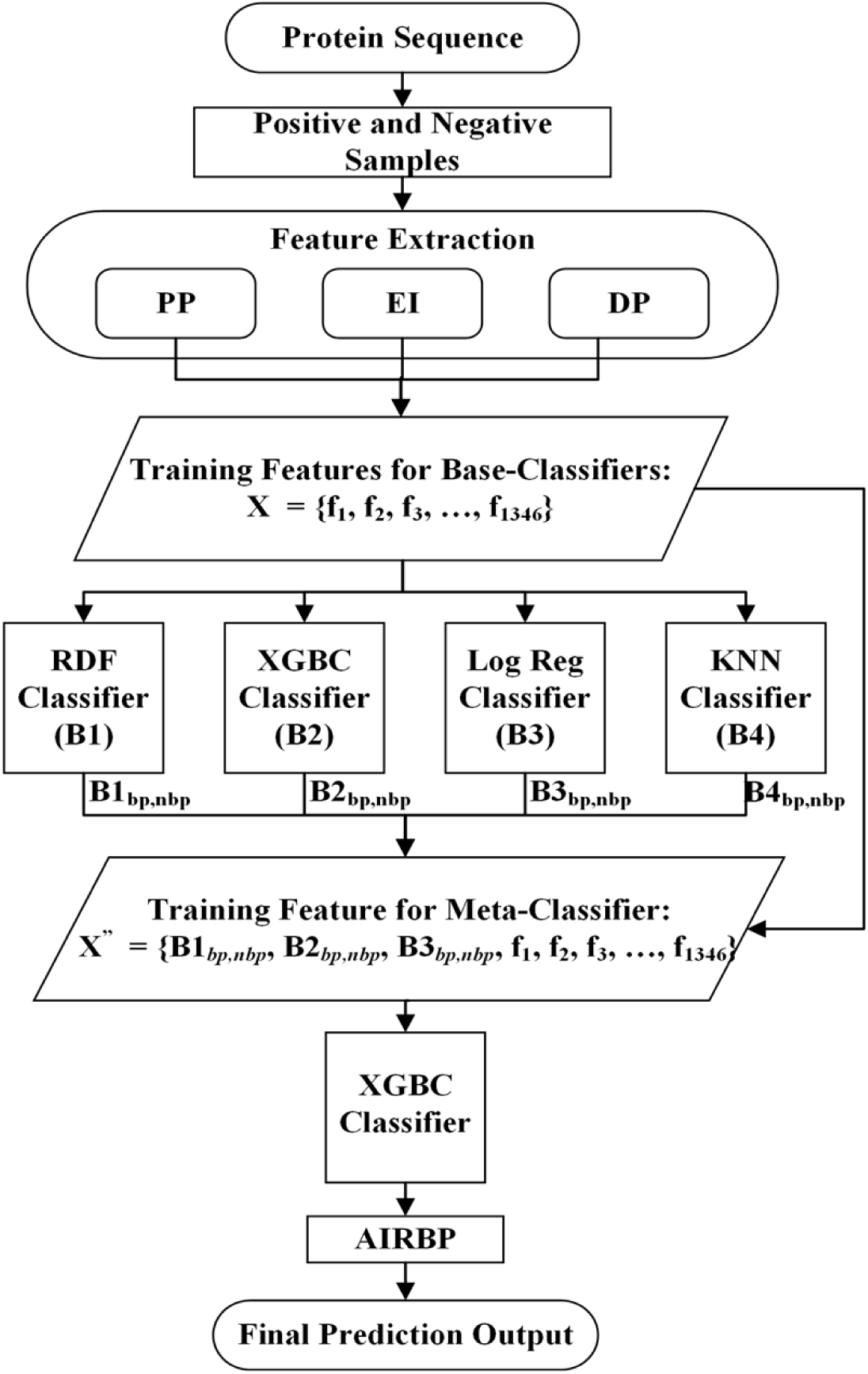
Illustrates the prediction framework of the AIRBP.

#### Features extracted from evolutionary information

In this section, we describe various feature extraction techniques utilized to obtain a fixed dimensional feature vector from the evolutionary information, called PSSM profile to encode protein sequence.

Evolutionary information is one of the most critical information useful for solving various biological problems and has been widely used in many research work (Iqbal, et al., 2015; Kumar, et al., 2007; Kumar, et al., 2008; Kumar, et al., 2011; Mishra, et al., 2018; Zhang and Liu, 2017). In this work, the evolutionary information in the form of the PSSM profile is directly obtained from the protein sequence and later transformed into a fixed dimensional vector. PSSM captures the conservation pattern in multiple alignments and preserves it as a matrix for each position in the alignment. The high score in the PSSM matrix indicates more conserved positions and the lower score indicates less conserved positions (Mishra, et al., 2018). For this study, we generated the PSSM profile for every protein sequence by executing three iterations of PSI-BLAST against NCBI’s non-redundant database (Altschul, et al., 1990). The evolutionary information in the PSSM profile is represented as a matrix of L*20 dimensions, where L is the length of the protein sequence. A particular element *M_i,j_* of the PSSM matrix represents the occurrence probability of the amino acid *i* at position *j* of a protein sequence.

##### PSSM-Distance transformation (PSSM-DT) features

We use two types of distance transformation techniques (Mishra, et al., 2018; Xu, et al., 2015): *i)* the PSSM distance transformation for same pairs of amino acids (PSSM-SDT); and *ii)* the PSSM distance transformation for different pairs of amino acids (PSSM-DDT), together known as PSSM-DT to extract fixed dimensional feature vectors of size 100 and 1900, respectively.

Utilizing PSSM-SDT, we compute the occurrence probabilities for the pairs of the same amino acids separated by a distance *D* along the sequence, which can be represented as:

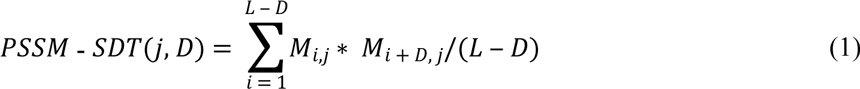

where *j* represents one type of the amino acid, *L* represents the length of the sequence, M*_i,j_* represents the PSSM score of amino acid *j* at position *i*, and *M_i+D,j_* represents the PSSM score of amino acid *j* at position *i+D.* Through this approach, 20 × *K* number of features were generated where *K* is the maximum range of *D* (*D =* 1,2, …, *K*).

Likewise, utilizing PSSM-DDT, we compute the occurrence probabilities for pairs of different amino acids separated by a distance *D* along the sequence, which can be represented as:

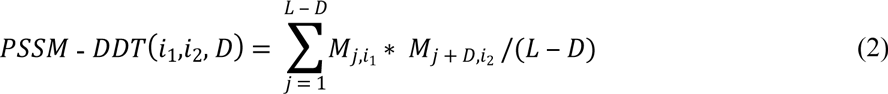

where, *i*_1_ and *i*_2_ represent two different types of amino acids. The total number of features obtained by PSSM-DDT is 380 × *K*. Here, we consider *K* = 5. Therefore a total of 100 features was obtained by PSSM-SDT and PSSM-DDT transformation techniques obtained a total of 1900 features.

##### Evolutionary distance transformation (EDT) features

Unlike PSSM-DT, the EDT approximately measures the non-co-occurrence probability of two amino acids separated by a specific distance *d* in a protein sequence from the PSSM profile (Mishra, et al., 2018; Zhang, et al., 2014). The EDT is calculated from the PSSM profile as:

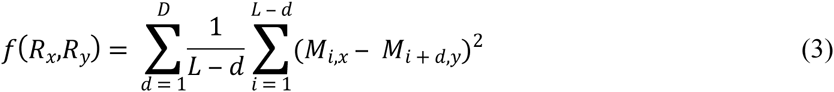

where *d* is the distance separating two amino acids, *D* is the maximum value of *d*, *M*_*i,x*_ and *M*_*i* + *d,y*_ are the elements in the PSSM profile, and *R*_*x*_ and *R*_*y*_ represent any of the 20 standard amino acids in the protein sequence. Here, the value of *D = L_min_-1* where *L_min_* is the length of the shortest protein sequence in the benchmark dataset. Using EDT, we obtain a feature vector of dimension 400.

#### Features extracted from disordered properties

In this section, we describe a feature extraction technique utilized to obtain a fixed dimensional feature vector from the residue-wise contact energy matrix to encode protein sequence.

RBPs are found to bind with RNA not only through classical structured RNA-binding domains but also through intrinsically disordered regions (IDRs) (Calabretta and Richard, 2015). For example, approximately 20% of the identified mammalian RBPs (∼170 proteins) were found to be disordered by over 80% (Järvelin, et al., 2016). The energy contribution of a large number of inter and intra-residual interactions in intrinsically disordered proteins (IDPs) cannot be approximated by the energy functions extracted from known structures (Hoque, et al., 2016; Iqbal, et al., 2015; Mishra and Hoque, 2017; Mishra, et al., 2016; Zhou and Skolnick, 2011) as IDPs lack a defined and ordered 3D structure (Babu, et al., 2011). Therefore, to inherently incorporate important information regarding the IDRs and amino acid interactions, we employed the predicted residue-wise contact energies (Dosztányi, et al., 2005) and molecular recognition features (MoRFs) (Sharma, et al., 2018), to encode the protein sequence.

##### Residue-wise contact energy matrix transformation (RCEM-T) features

We adopted the predicted residue-wise contact energy matrix (RCEM) derived in (Dosztányi, et al., 2005), by the least square fitting of 674 proteins primary sequence with the contact energies derived from the tertiary structure of 785 proteins. As shown in Table 1, the RCEM is a 20 × 20 dimensional matrix that contains residue-wise contact energy for 20 standard amino acids. For a protein sequence of length *L*, an *L* × 20 dimensional matrix *M* is obtained which holds a 20-dimensional vector for each amino acid in a protein sequence. The resulting matrix *M* is then encoded into a feature vector of 20 dimensional by computing the column-wise sum as:

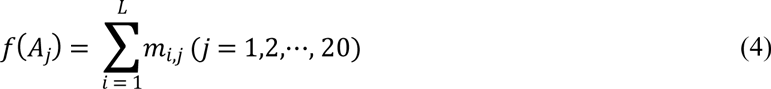

where *m_i,j_* is the element of matrix *M*, *i* is the amino acid index in a sequence, and *j* represents 20 standard amino acid types. The final feature vector, *RCEM* − *T* = [*v*_1_, *v*_2_, ⋯, *v*_20_] is obtained by dividing each element in *RCEM-T* by the sum of all the elements in the same vector. Considering *V_s_* as the sum of all the elements in the RCEM-T vector, each component of the final *RCEM-T* vector can be represented as:

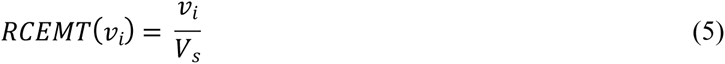

**Table 1.** RCEM table used to obtain RCEM-T features.

|  | A | C | D | E | F | G | H | I | K | L | M | N | P | Q | R | S | T | V | W | Y |
| --- | --- | --- | --- | --- | --- | --- | --- | --- | --- | --- | --- | --- | --- | --- | --- | --- | --- | --- | --- | --- |
| A | -1.65 | -2.83 | 1.16 | 1.8 | -3.73 | -0.41 | 1.9 | -3.69 | 0.49 | -3.01 | -2.08 | 0.66 | 1.54 | 1.2 | 0.98 | -0.08 | 0.46 | -2.31 | 0.32 | -4.62 |
| C | -2.83 | -39.58 | -0.82 | -0.53 | -3.07 | -2.96 | -4.98 | 0.34 | -1.38 | -2.15 | 1.43 | -4.18 | -2.13 | -2.91 | -0.41 | -2.33 | -1.84 | -0.16 | 4.26 | -4.46 |
| D | 1.16 | -0.82 | 0.84 | 1.97 | -0.92 | 0.88 | -1.07 | 0.68 | -1.93 | 0.23 | 0.61 | 0.32 | 3.31 | 2.67 | -2.02 | 0.91 | -0.65 | 0.94 | -0.71 | 0.90 |
| E | 1.8 | -0.53 | 1.97 | 1.45 | 0.94 | 1.31 | 0.61 | 1.3 | -2.51 | 1.14 | 2.53 | 0.2 | 1.44 | 0.1 | -3.13 | 0.81 | 1.54 | 0.12 | -1.07 | 1.29 |
| F | -3.73 | -3.07 | -0.92 | 0.94 | -11.25 | 0.35 | -3.57 | -5.88 | -0.82 | -8.59 | -5.34 | 0.73 | 0.32 | 0.77 | -0.4 | -2.22 | 0.11 | -7.05 | -7.09 | -8.80 |
| G | -0.41 | -2.96 | 0.88 | 1.31 | 0.35 | -0.2 | 1.09 | -0.65 | -0.16 | -0.55 | -0.52 | -0.32 | 2.25 | 1.11 | 0.84 | 0.71 | 0.59 | -0.38 | 1.69 | -1.90 |
| H | 1.9 | -4.98 | -1.07 | 0.61 | -3.57 | 1.09 | 1.97 | -0.71 | 2.89 | -0.86 | -0.75 | 1.84 | 0.35 | 2.64 | 2.05 | 0.82 | -0.01 | 0.27 | -7.58 | -3.20 |
| I | -3.69 | 0.34 | 0.68 | 1.3 | -5.88 | -0.65 | -0.71 | -6.74 | -0.01 | -9.01 | -3.62 | -0.07 | 0.12 | -0.18 | 0.19 | -0.15 | 0.63 | -6.54 | -3.78 | -5.26 |
| K | 0.49 | -1.38 | -1.93 | -2.51 | -0.82 | -0.16 | 2.89 | -0.01 | 1.24 | 0.49 | 1.61 | 1.12 | 0.51 | 0.43 | 2.34 | 0.19 | -1.11 | 0.19 | 0.02 | -1.19 |
| L | -3.01 | -2.15 | 0.23 | 1.14 | -8.59 | -0.55 | -0.86 | -9.01 | 0.49 | -6.37 | -2.88 | 0.97 | 1.81 | -0.58 | -0.6 | -0.41 | 0.72 | -5.43 | -8.31 | -4.90 |
| M | -2.08 | 1.43 | 0.61 | 2.53 | -5.34 | -0.52 | -0.75 | -3.62 | 1.61 | -2.88 | -6.49 | 0.21 | 0.75 | 1.9 | 2.09 | 1.39 | 0.63 | -2.59 | -6.88 | -9.73 |
| N | 0.66 | -4.18 | 0.32 | 0.2 | 0.73 | -0.32 | 1.84 | -0.07 | 1.12 | 0.97 | 0.21 | 0.61 | 1.15 | 1.28 | 1.08 | 0.29 | 0.46 | 0.93 | -0.74 | 0.93 |
| P | 1.54 | -2.13 | 3.31 | 1.44 | 0.32 | 2.25 | 0.35 | 0.12 | 0.51 | 1.81 | 0.75 | 1.15 | -0.42 | 2.97 | 1.06 | 1.12 | 1.65 | 0.38 | -2.06 | -2.09 |
| Q | 1.2 | -2.91 | 2.67 | 0.1 | 0.77 | 1.11 | 2.64 | -0.18 | 0.43 | -0.58 | 1.9 | 1.28 | 2.97 | -1.54 | 0.91 | 0.85 | -0.07 | -1.91 | -0.76 | 0.01 |
| R | 0.98 | -0.41 | -2.02 | -3.13 | -0.4 | 0.84 | 2.05 | 0.19 | 2.34 | -0.6 | 2.09 | 1.08 | 1.06 | 0.91 | 0.21 | 0.95 | 0.98 | 0.08 | -5.89 | 0.36 |
| S | -0.08 | -2.33 | 0.91 | 0.81 | -2.22 | 0.71 | 0.82 | -0.15 | 0.19 | -0.41 | 1.39 | 0.29 | 1.12 | 0.85 | 0.95 | -0.48 | -0.06 | 0.13 | -3.03 | -0.82 |
| T | 0.46 | -1.84 | -0.65 | 1.54 | 0.11 | 0.59 | -0.01 | 0.63 | -1.11 | 0.72 | 0.63 | 0.46 | 1.65 | -0.07 | 0.98 | -0.06 | -0.96 | 1.14 | -0.65 | -0.37 |
| V | -2.31 | -0.16 | 0.94 | 0.12 | -7.05 | -0.38 | 0.27 | -6.54 | 0.19 | -5.43 | -2.59 | 0.93 | 0.38 | -1.91 | 0.08 | 0.13 | 1.14 | -4.82 | -2.13 | -3.59 |
| W | 0.32 | 4.26 | -0.71 | -1.07 | -7.09 | 1.69 | -7.58 | -3.78 | 0.02 | -8.31 | -6.88 | -0.74 | -2.06 | -0.76 | -5.89 | -3.03 | -0.65 | -2.13 | -1.73 | -12.39 |
| Y | -4.62 | -4.46 | 0.9 | 1.29 | -8.8 | -1.9 | -3.2 | -5.26 | -1.19 | -4.9 | -9.73 | 0.93 | -2.09 | 0.01 | 0.36 | -0.82 | -0.37 | -3.59 | -12.39 | -2.68 |

##### Molecular recognition features (MoRFs)

MoRFs, also sometimes known as molecular recognition elements (MoREs), are disordered regions in a protein that exhibit various molecular recognition and binding functions (Vacic, et al., 2007). Post-translational modifications (PTMs) can induce disorder to order transitions of IDPs upon binding with their binding partners which could be either RNA, DNA, proteins, lipids, carbohydrates or other small molecules (Bah and Forman-Kay, 2016; Lina, et al., 2017). MoRFs play a vital role in various biological functions of IDPs located within long disordered protein sequences (Mohan, et al., 2006; Sharma, et al., 2018; Sharma, et al., 2018; Sharma, et al., 2018). Additionally, Mohan *et al*. suggest that functionally significant residual structures exist in MoRF regions prior to the actual binding (Mohan, et al., 2006). These residual structures could, therefore, be useful in the prediction of binding between proteins and RNA. Here, to capture the functional properties of IDRs that may bind to RNAs, we employ a single predicted MoRFs score as a feature. To obtain a single predicted MoRFs score, first, the residue-wise predicted MoRFs scores are obtained from the OPAL program (Sharma, et al., 2018). Then, a single predicted MoRFs score is computed by taking a ratio of the sum of the residue-wise MoRFs score and the length of the protein sequence.

### Performance evaluation

To evaluate the performance of AIRBP, we adopted a widely used 10-fold CV and the independent testing approach. In the process of 10-fold CV, the dataset is segmented into 10 parts, which are each of about the same size. When one fold is kept aside for testing, the remaining 9 folds are used to train the classifier. This process of training and test is repeated until each fold has been kept aside once for testing and consequently, the test accuracies of each fold are combined to compute the average (Hastie, et al., 2009). Unlike a 10-fold CV, in independent testing, the classifier is trained with the benchmark dataset and consequently tested using the independent test dataset. While independent testing, it is ensured that none of the samples in the independent test set are present in the benchmark dataset. We used several performance evaluation metrics listed in Table 2 as well as ROC and AUC to test the performance of the proposed method as well as to compare it with the existing approaches. AUC is the area under the receiver operating characteristics (ROC) curve which is used to evaluate how well a predictor separates two classes of information (RNA-binding and non-binding protein).

**Table 2.**
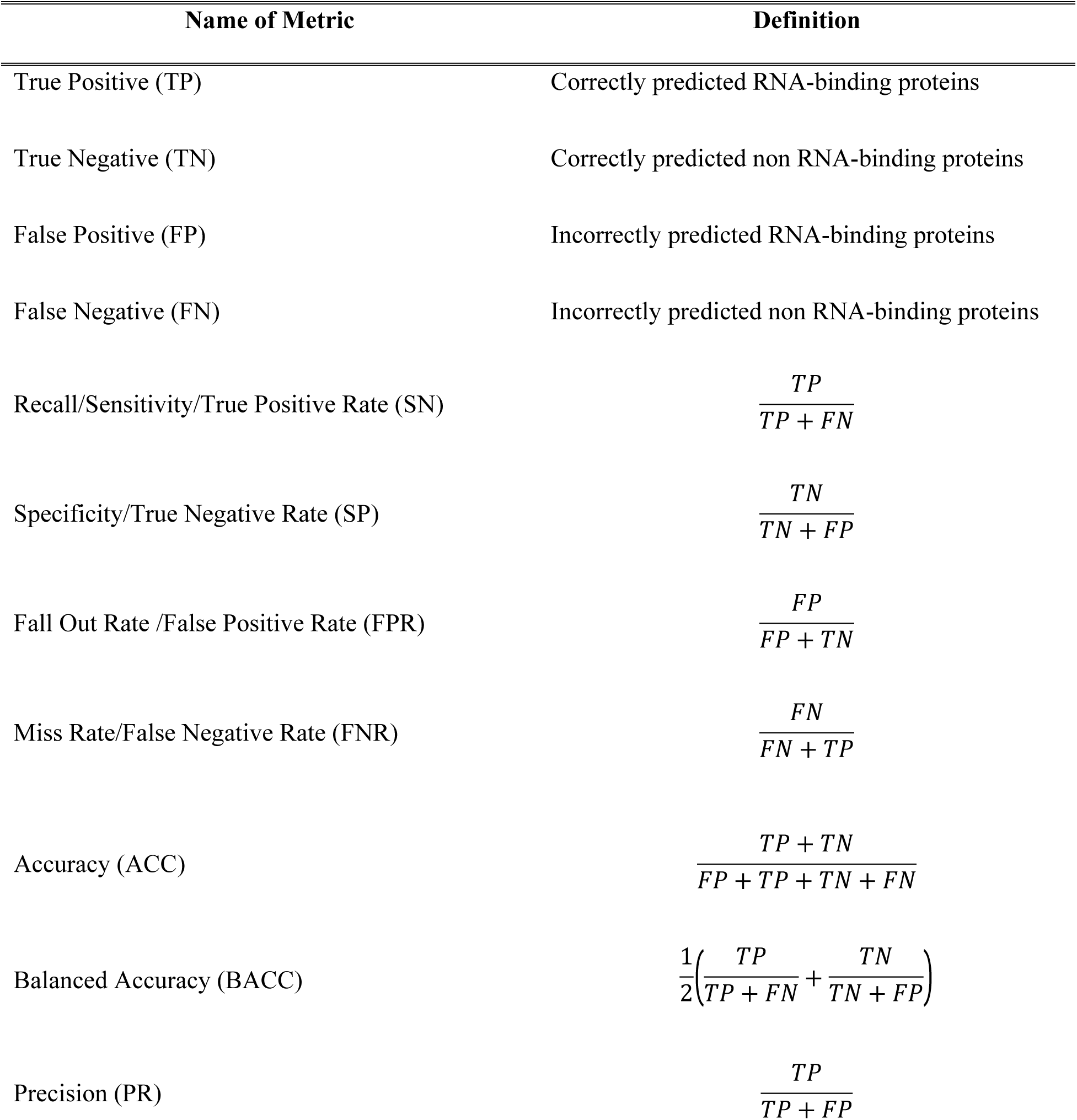
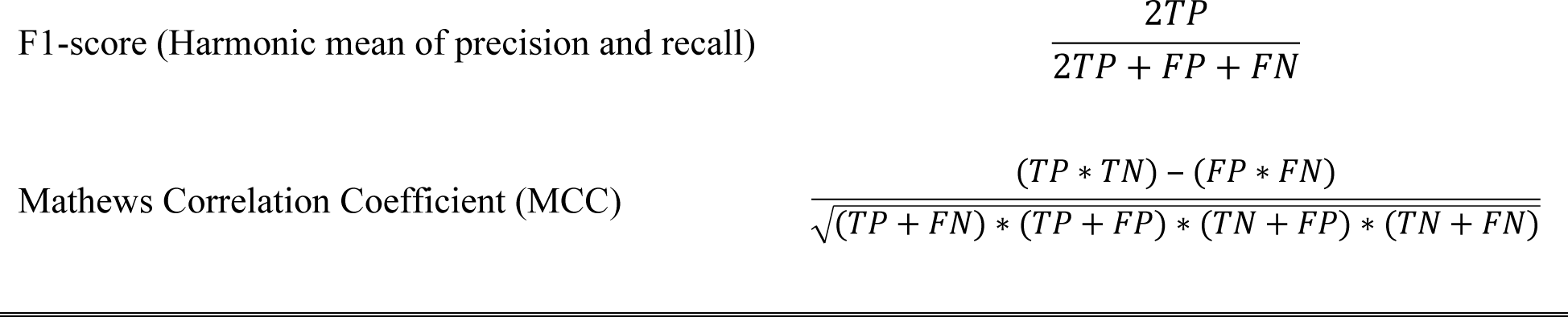
Name and definition of the evaluation metric.

### Feature Selection

During the feature extraction process, we collected a feature vector of 2603 dimensions, which is significantly large. Therefore, to reduce the feature space and select the relevant features that could help improve the classification accuracy, we adopted two distinct feature selection approaches, namely Incremental Feature Selection (IFS) and Genetic Algorithm (GA), a class of evolutionary algorithm, based feature selection. A detailed description of the two feature selection approaches is provided below.

#### Feature Selection using IFS

IFS starts with an empty feature vector and a feature group is added to the feature vector if the addition of the feature group to the feature vector improves the performance of the predictor. In case, by adding the new feature group, the accuracy of the predictor is reduced, this feature group is discarded, and a new feature group is tested iteratively. During IFS, we performed a 10-fold CV on the benchmark dataset using XGBoost as a predictor. The values of XGBoost parameters: max_depth, eta, silent, objective, num_class, n_estimators, min_child_weight, subsample, scale_pos_weight, tree_method and max_bin were set to 6, 0.1, 1, ‘multi:softprob’, 2, 100, 5, 0.9, 3, ‘hist’ and 500, respectively and the rest of the parameters were set to their default value. We used ACC as the evaluation metric to decide whether the new feature group will a to the feature vector or not. In our implementation of IFS, only the Vander Waals Volume feature group was ignored from the feature vector as the addition of this feature decreased the ACC of the predictor. Therefore, through IFS, 2582 features out of 2603 features were selected as relevant features.

#### Feature Selection using GA

GA is a population-based stochastic search technique that mimics the natural process of evolution. It contains a population of chromosomes where each chromosome represents a possible solution to the problem under consideration. In general, a GA operates by initializing the population randomly, and by iteratively updating the population through various operators including elitism, crossover, and mutation to discover, prioritize and recombine good building blocks present in parent chromosomes to finally obtain fitter chromosome (Hoque, et al., 2010; Hoque, et al., 2007; Hoque and Iqbal, 2017).

Encoding the solution of the problem under consideration in the form of chromosomes and computing the fitness of the chromosomes are two important steps in setting up the GA. Here, to perform feature selection, we encode each feature *f*_*i*_ in our feature space *F* = [*f*_1_, *f*_2_, ⋯, *f*_*n*_] by a single bit of 1/0 in a chromosome space where the value of 1 represents that the *i*^th^ feature is selected, and the value of 0 represents that the *i*^th^ feature is not selected. The length of the chromosome space is equal to the length of the feature space. Moreover, to compute the fitness of the chromosome, we use the XGBoost algorithm (Chen and Guestrin, 2016). The choice of XGBoost was made because of its fast execution time and reasonable performance compared to other machine learning classifiers. During feature selection, the values of XGBoost parameters: max_depth, eta, silent, objective, num_class, n_estimators, min_child_weight, subsample, scale_pos_weight, tree_method, and max_bin were set to 6, 0.1, 1, ‘multi:softprob’, 2, 100, 5, 0.9, 3, ‘hist’ and 500, respectively and the rest of the parameters were set to their default value. The values of the XGBoost parameters mentioned above were identified through the hit and trial approach. In our implementation, the objective fitness is defined as:

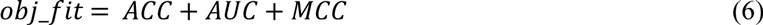

where, ACC is the accuracy, AUC is the area under the receiver operating characteristic curve and MCC is the Matthews Correlation Coefficient. To evaluate the fitness of the chromosome, a new data space *D* is obtained which only includes the features for which the chromosome bit is 1. The values of ACC, AUC and MCC metrics of the *obj_fit* are obtained by performing a 10-fold CV on a new data space *D* using the XGBoost algorithm. Furthermore, the additional parameters of the GA in our implementation were set to a population size of 20, maximum generation to 300, elite-rate to 5%, crossover-rate to 90%, and mutation rate to 50%. Through this GA based feature selection, only 1346 features out of 2603 features were selected as relevant features. Therefore, we were able to achieve two-fold benefits from the GA based features selection which are significantly reduced feature space and relevant features. Finally, we noticed that at least one of the features from each type of feature we extracted was present in the feature set selected by GA. Therefore, all the feature types extracted in this study were found to be essential for the prediction of RBPs.

### Framework of AIRBP

To develop AIRBP predictor for RBPs prediction, we adopted an idea of stacking based machine learning approach (Wolpert, 1992) which, has recently been successfully applied to solve various bioinformatics problems (Hu, et al., 2015; Iqbal and Hoque, 2018; Mishra, et al., 2018; Nagi and Bhattacharyya, 2013). Stacking is an ensemble-based machine learning approach, which collects information from multiple models in different phases and combines them to form a new model. Stacking is considered to yield more accurate results than the individual machine learning methods as the information gained from more than one predictive model minimizes the generalization error. The stacking framework includes two-levels of classifiers, where the classifiers of the first-level are called base-classifiers and the classifiers of the second-level are called meta-classifiers. In the first level, a set of base-classifiers C_1_, C_2_, …, C_N_ is employed (Džeroski and Ženko, 2004). The prediction probabilities from the base-classifiers are combined using a meta-classifier to reduce the generalization error and improve the accuracy of the predictor. To enrich the meta-classifier with necessary information on the problem space, the classifiers at the base-level are selected such that their underlying operating principle is different from one another (Mishra, et al., 2018; Nagi and Bhattacharyya, 2013).

To select the classifiers to use in the first and second level of the AIRBP stacking framework, we analyzed the performance of six individual classification methods: *i)* Random Decision Forest (RDF) (Ho, 1995); *ii)* Bagging (Bag) (Breiman, 1996); *iii)* Extra Tree (ET) (Geurts, et al., 2006); iv) Extreme Gradient Boosting (XGBoost or XGB) (Chen and Guestrin, 2016); v) Logistic Regression (LogReg) (Hastie, et al., 2009; Szilágyi and Skolnick, 2006); and vi) K-Nearest Neighbor (KNN) (Altman, 1992). The algorithms and their configuration details are briefly discussed below.

i. *RDF:* RDF (Ho, 1995) constructs a multitude of decision trees, each of which is trained on a random subset of the training data. The sub-set used to create a decision tree is constructed from a given set of observations of training data by taking ‘m’ observations at random and with replacement (a.k.a. Bootstrap Sampling). Next, the final predictions are achieved by aggregating the prediction from the individual decision trees. For classification, the final prediction is made by computing the mode (the value that appears most often) of the classes (in our case: whether a protein is RNA-binding or non-binding). In our implementation of the RDF, we used bootstrap samples to construct 1,000 trees (n_estimators=1,000) in the forest, and the rest of the parameters were set to their default value.
ii. *Bag:* Bag (Breiman, 1996) machine learning algorithm operates by forming a class of algorithms that creates several instances of a base classifier/estimator on random subsets of the training samples and consequently combines their individual predictions to yield a final prediction. It reduces the variance in the prediction. In our study, the BAG classifier was fit on multiple subsets of data using Bootstrap Sampling using 1,000 decision trees (n_estimators=1,000) and the rest of the parameters were set to their default value.
iii. *ET:* Extremely randomized tree (ET) classifier (Geurts, et al., 2006) operates by fitting several randomized decision trees (a.k.a. extra-trees) on various sub-sets and uses averaging to improve the prediction accuracy and control over-fitting. In our implementation, the ETC model was constructed using 1,000 trees (n_estimators=1,000) and the quality of a split was assessed by the Gini impurity index. The rest of the parameters were set to their default value.
iv. *XGBoost:* XGBoost (Chen and Guestrin, 2016) follows the same principle of gradient boosting as the Gradient Boosting Classifier (GBC). GBC (Friedman, 2001) involves three elements: (a) a loss function to be optimized, (b) a weak learner to make predictions, and (c) an additive model to add weak learners to minimize the loss function. The objective of GBC is to minimize the loss of the model by adding weak learners in a stage-wise fashion using a procedure similar to gradient descent. The existing weak learners in the model are remained unchanged while adding new weak learners. The output from the new learner is added to the output of the existing sequence of learners to correct or improve the final output of the model. Unlike GBC, XGBoost performs more regularized model formalization to control over-fitting, which results in better performance. In addition to increased performance, XGBoost provides higher computational speed. In our configuration of XGBoost, the values of parameters: colsample_bytree, gamma, min_child_weight, learning_rate, max_depth, n_estimators, and subsample ratio were optimized to achieve the best 10-fold cross-validation accuracy using a grid search (Bergstra and Bengio, 2012) technique. The best values of the parameters: colsample_bytree, gamma, min_child_weight, learning_rate, max_depth, n_estimators, and subsample ratio were found to be 0.6, 0.3, 1.5, 0.07, 5, 10000 and 0.95, respectively. And, the rest of the parameters were set to their default value.
v. *LogReg:* LogReg (a.k.a. logit or MaxEnt) (Hastie, et al., 2009; Szilágyi and Skolnick, 2006) is a machine learning classifier that measures the relationship between the dependent categorical variable (in our case: a RNA-binding or non-binding proteins) and one or more independent variables by generating an estimation probability using logistic regression. In our implementation, we set all the parameters of LogReg to their default values.
vi. *KNN:* KNN (Altman, 1992) is a non-parametric and lazy learning algorithm. Non-parametric means it does not make any assumption for underlying data distribution, rather it creates models directly from the dataset. Furthermore, lazy learning means it does not need any training data points for a model generation rather uses the training data while testing. It works by learning from the K number of training samples closest in the distance to the target point in the feature space. The classification decision is made based on the majority-votes obtained from the K nearest neighbors. Here, we set the value of K to 9 and the rest of the parameters to their default value.

All the classification methods mentioned above are built and optimized using python’s Scikit-learn library (Pedregosa, et al., 2012). In order to design a stacking framework for AIRBP, we evaluated the different combinations of base-classifiers and finally selected the one that provided the highest performance.

The set of stacking framework tested are:

1. SF1: RDF, XGBoost, LogReg, KNN in base-level and XGBoost in meta-level,
2. SF2: Bag, XGBoost, LogReg, KNN in base-level and XGBoost in meta-level and
3. SF3: ET, XGBoost, LogReg, KNN in base-level and XGBoost in meta-level.

Here, the choice of base-level classifiers is made such that the underlying principle of learning of each of the classifiers is different from each other (Mishra, et al., 2018). For example, in SF1, SF2 and SF3 the tree-based classifiers RDF, Bag and ET are individually combined with the other two methods LogReg and KNN to learn different information from the problem-space. Additionally, for each of the combination SF1, SF2 and SF3, the XGBoost classifier is used both in the base as well as in the meta-level because it performed best among all the other individual methods applied in this work. While examining the 10-fold CVs performance of the above three combinations, we found that the first stacking framework, SF1 attains the highest performance. Therefore, we employ four classifiers RDF, XGBoost, LogReg, and KNN as the base classifiers and another XGBoost as the meta-classifier in AIRBP stacking framework. In AIRBP, the probabilities of both the classes (RBP and non-RBP) generated by the four base-classifiers are combined with the 1346 features selected by GA and provided as an input features to the meta-classifier which eventually provides the prediction for RBPs.

## Results

In this section, we first demonstrate the results of the feature selection. Then, we show the performance comparison of potential base-classifiers and stacking frameworks. Finally, we report the performance of AIRBP on the benchmark dataset and three independent test datasets and consequently compare it with the existing methods

### Feature Selection

To reduce the feature space and select the relevant features that support the classification accuracy, we adopted the IFS and GA based feature selection approach. Through IFS and GA, 2582 and 1346 features out of 2603 total features were selected as relevant features, respectively. From Table 4, we observe that IFS could not reduce the feature space as significantly as GA. Additionally, the performance of XGBoost after IFS did not improve by significant value and is lower than the performance resulted from the GA-based feature selection. We found that the benefit of GA feature selection was two folds, considerable reduction of feature space and identification of relevant features along with improved performance. To assess the impact of individual feature groups that are obtained from GA, we performed feature contribution analysis using the XGBoost classifier. The details of the feature contribution analysis are provided below.

**Table 3.**
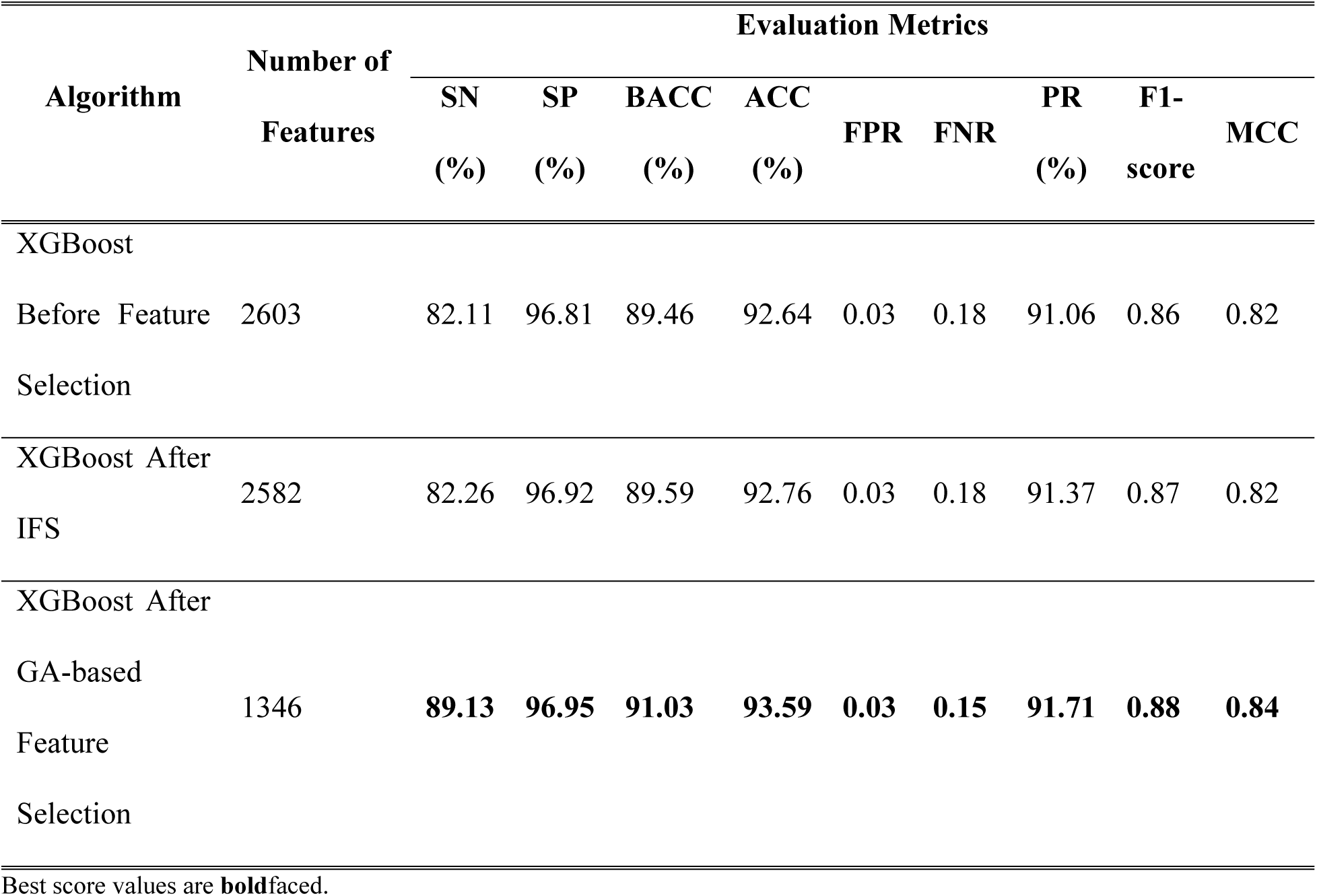
Comparison of RBPs prediction results on benchmark dataset before and after feature selection.

**Table 4.** Contribution of features on the performance of the XGBoost classifier obtained through 10-fold cross-validation on the benchmark dataset.

| Features | SN<br>(%) | SP<br>(%) | ACC<br>(%) | BACC<br>(%) | fpr | fnr | PRE<br>(%) | F1 | MCC |
| --- | --- | --- | --- | --- | --- | --- | --- | --- | --- |
| EDT | 65.78 | 93.17 | 85.40 | 79.47 | 0.07 | 0.34 | 79.23 | 0.72 | 0.63 |
| EDT+SDT | 74.09 | 95.05 | 89.10 | 84.57 | 0.05 | 0.26 | 85.56 | 0.79 | 0.72 |
| EDT+SDT+DDT | 78.35 | 96.18 | 91.12 | 87.27 | 0.04 | 0.22 | 89.04 | 0.83 | 0.78 |
| EDT+SDT+DDT+Hydrophobicity | 79.29 | 96.81 | 91.84 | 88.05 | 0.03 | 0.21 | 90.77 | 0.85 | 0.79 |
| EDT+SDT+DDT+Hydrophobicity +Polarity | 86.23 | 97.27 | 94.14 | 91.75 | 0.03 | 0.14 | 92.59 | 0.89 | 0.85 |
| EDT+SDT+DDT+Hydrophobicity +Polarity+Polarizability | 86.95 | 96.82 | 94.02 | 91.88 | 0.03 | 0.13 | 91.56 | 0.89 | 0.85 |
| EDT+SDT+DDT+Hydrophobicity<br>+Polarity+Polarizability+Van | 87.28 | 97.02 | 94.26 | 92.15 | 0.03 | 0.13 | 92.07 | 0.90 | 0.86 |
| EDT+SDT+DDT+Hydrophobicity<br>+Polarity+Polarizability+Van+CT | 87.53 | 97.16 | 94.43 | 92.35 | 0.03 | 0.12 | 92.44 | 0.90 | 0.86 |
| EDT+SDT+DDT+Hydrophobicity<br>+Polarity+Polarizability+Van+CT+ACC | 87.97 | 97.28 | 94.62 | 92.62 | 0.03 | 0.12 | 92.76 | 0.90 | 0.87 |
| EDT+SDT+DDT+Hydrophobicity<br>+Polarity+Polarizability+Van+CT+ACC+SS | 88.40 | 97.47 | 94.90 | 92.93 | 0.03 | 0.12 | 93.25 | 0.91 | 0.87 |
| EDT+SDT+DDT+Hydrophobicity<br>+Polarity+Polarizability+Van+CT+ACC+SS+RCEM | 89.12 | 97.37 | 95.03 | 93.24 | 0.03 | 0.11 | 93.06 | 0.91 | 0.88 |
| EDT+SDT+DDT+Hydrophobicity<br>+Polarity+Polarizability+Van+CT+ACC+SS+RCEM+MoRFs | 89.01 | 97.35 | 94.99 | 93.18 | 0.03 | 0.11 | 93.01 | 0.91 | 0.88 |

During feature contribution analysis, the values of XGBoost parameters: colsample_bytree, gamma, min_child_weight, learning_rate, max_depth, n_estimators, subsample were set to 0.6, 0.3, 1.5, 0.07, 5, 10000, and 0.95, respectively and the rest of the parameters were set to their default value. The values of the XGBoost parameters mentioned above were identified through the grid search approach. Table 4 shows the impact of the addition of individual features on the performance of the XGBoost classifier. Starting with the first feature group, EDT, we build several XGBoost classifiers by adding one feature group in the feature vector at a time, through 10-fold cross-validation on the benchmark dataset. Table 4 shows the results of feature contribution analysis.

From Table 4, we can see that incrementally adding the feature group into the feature vector improves the performance of the XGBoost classifier obtained through 10-fold cross-validation on the benchmark dataset. The improvement in the performance of the XGBoost classifier obtained by the sequential addition of feature group into feature vector indicates that all the features implemented in our study are useful. Notably, we can observe that the addition of SDT and DDT features itself improved the MCC from 0.63 to 0.72 and 0.78, respectively. Further, we can also observe that the addition of the RCEM and MoRFs feature improves the MCC of the predictor from 0.87 to 0.88. This indicates that the addition of RCEM and MoRFs features alone provides an improvement of 1.15%.

### Selection of Classifiers for Stacking

To select the methods to use as the base and the meta-classifiers, we analyzed the performance of six different machine learning algorithms: RDF, Bag, ET, XGBoost, LogReg, and KNN on the benchmark dataset through 10-fold CV approach. The performance comparison of the individual classifiers on the benchmark dataset is shown in Table 5.

**Table 5.**
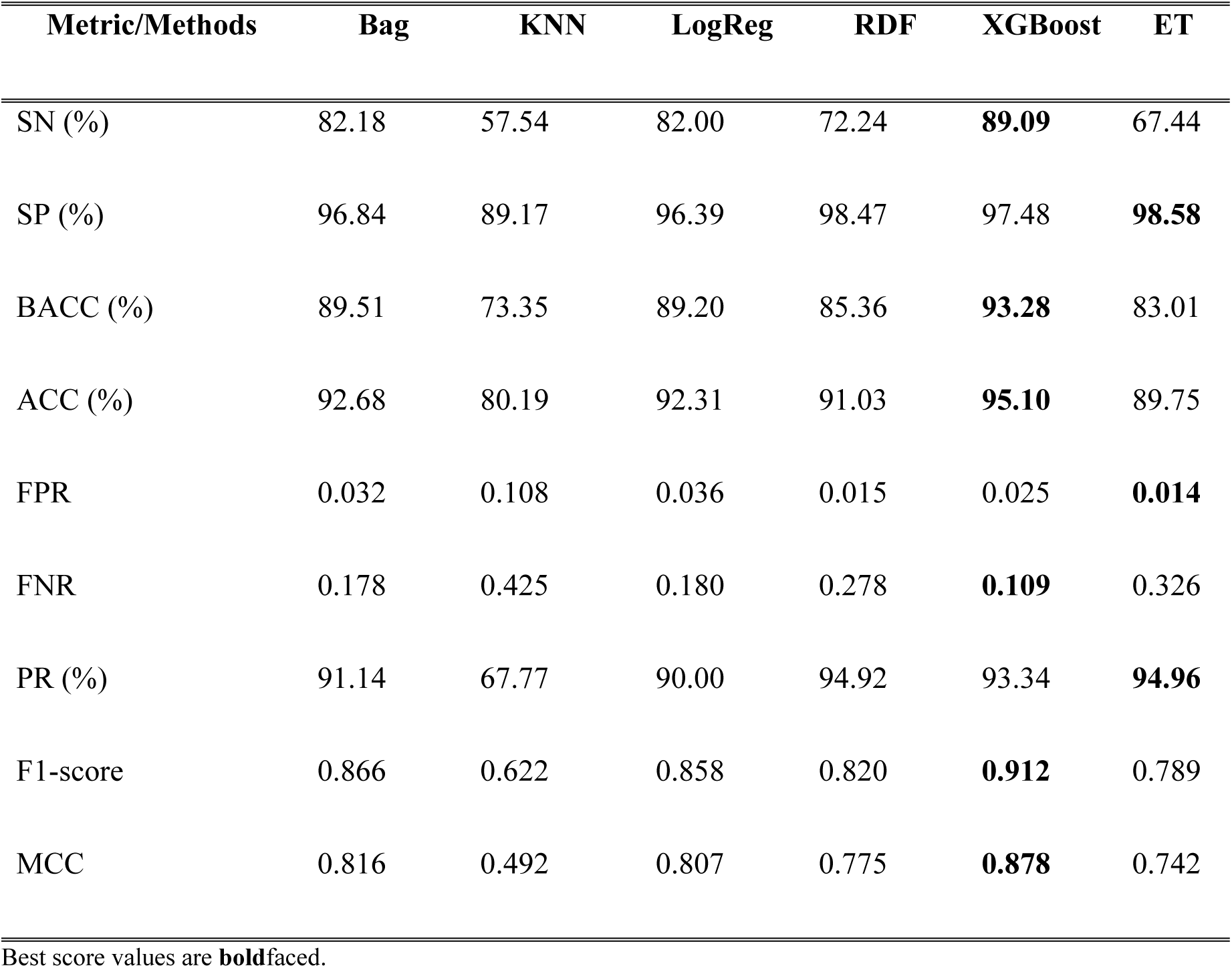
Comparison of various machine learning algorithms on the benchmark dataset through a 10-fold CV.

Table 5 further shows that the optimized XGBoost is the best performing classifier among six different classifiers implemented in our study, in terms of sensitivity, balanced accuracy, accuracy, FNR, F1-score, and MCC. Moreover, the optimized XGBoost attains sensitivity, balanced accuracy, accuracy, FNR, F1-score, and MCC of 89.09%, 93.28%, 0.109, 0.912, and 0.878, respectively. Besides, the ET classifier attains the highest specificity, FPR, and precision of 98.58%, 0.014, and 94.96%, respectively. As the benchmark dataset is highly imbalanced, we consider MCC as the deciding score as it provides the balanced measure of any predictor trained on an imbalanced dataset. Furthermore, it is evident from Table 5 that the MCC of the optimized XGBoost is 18.33%, 13.29%, 8.79%, 78.46%, and 7.59% higher than ET, RDF, LogReg, KNN, and Bag, respectively. The greater performance of the XGBoost algorithm motivated us to use it both as a base as well as a meta-classier in the AIRBP prediction framework.

To further select the classifiers to be used at the base-level, we adopted the guidelines of base-classifier selection based on different underlying principles. Therefore, we used KNN and LogReg as two additional classifiers at the base-level. Then, we added a single tree-based ensemble method out of three methods, RDF, Bag, and ET, at a time as the fourth base-classifier and designed three different combinations of stacking framework, namely SF1, SF2, and SF3. The performance comparison of SF1, SF2 and SF3 stacking framework on the benchmark dataset using 10-fold CV are presented in Table 6. Table 6 demonstrates that both SF1 and SF3 outperform SF2. Moreover, SF1 gained similar performance compared to SF3 in terms of ACC, F1-score, and MCC. Since our aim through this research is to build a robust system that makes correct predictions, we choose precision and FPR to be our deciding metrics. From Table 6, it is evident that SF1 has higher precision and lower FPR compared to SF2. Hence, we select SF1, which includes RDF, XGBoost, LogReg, and KNN as base-classifiers and another XGBoost as a meta-classifier, as our final predictor.

**Table 6.** Comparison of different stacking framework with a different set of base-classifiers on the benchmark dataset through a 10-fold CV.

| Metric/Methods | SF1 | SF2 | SF3 |
| --- | --- | --- | --- |
| SN (%) | **82.18** | 57.54 | 82.00 |
| SP (%) | **96.84** | 89.17 | 96.39 |
| BACC (%) | **89.51** | 73.35 | 89.20 |
| ACC (%) | **92.68** | 80.19 | 92.31 |
| FPR | **0.032** | 0.108 | 0.036 |
| FNR | **0.178** | 0.425 | 0.180 |
| PR (%) | **91.14** | 67.77 | 90.00 |
| F1-score | **0.866** | 0.622 | 0.858 |
| MCC | **0.816** | 0.492 | 0.807 |

Best score values are **boldfaced**.

#### Performance Comparison with Existing Approaches on the Benchmark Dataset

Here, we compare the performance of AIRBP with RBPPred (Zhang and Liu, 2017) on the benchmark dataset using the 10-fold CV approach. RBPPred is a top-performing existing approach for the prediction of RBPs directly from the sequence. Furthermore, it is to be noted that AIRBP uses the same benchmark dataset as RBPPred. Therefore, for the comparison, the quantities for all the evaluation metrics for RBPPred are obtained from Zhang and Liu (Zhang and Liu, 2017). The prediction results of AIRBP and RBPPred on benchmark dataset computed using 10-fold CV are listed in Table 7.

**Table 7.**
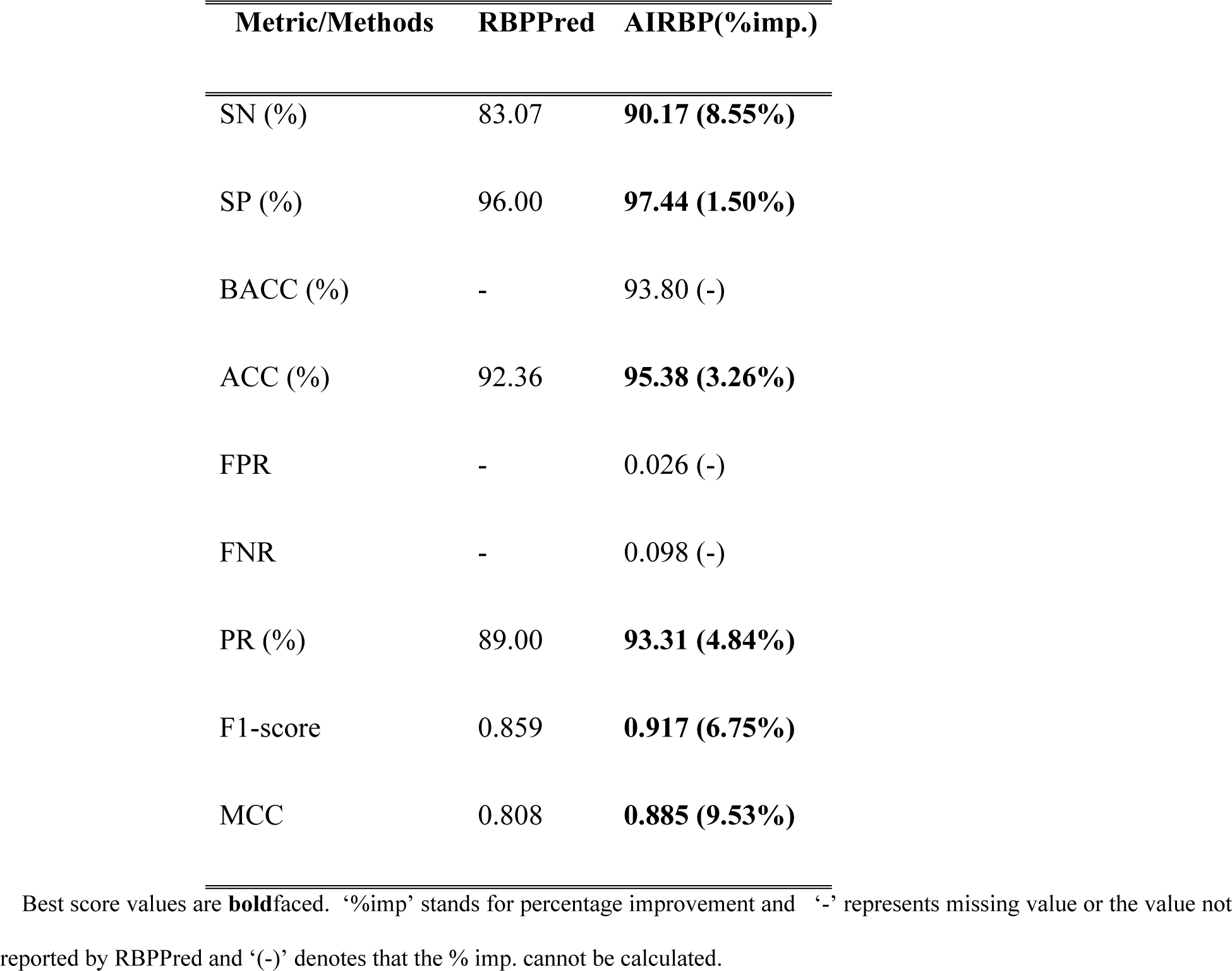
Comparison of AIRBP with existing method on benchmark dataset through 10-fold CV

From Table 7, we observed that AIRBP outperforms RBPPred based on all the evaluation metrics applied in this study. Particularly, AIRBP provides 8.55%, 1.50%, 3.26%, 4.84%, 6.75% and 9.53% improvement over RBPPred based on SN, SP, ACC, PR, F1-score and MCC, respectively. Besides, in Table 7, we report the values of BACC, FPR, and FNR only for the AIRBP predictor as the values of these metrics were not reported by RBPPred. Since our benchmark dataset is highly imbalanced (contains 2767 RBPs and 6987 non-RBPs), which also reflects the natural frequency, we focus on comparing the predictors based on MCC and F1-score. MCC considers true and false positives as well as negatives and is generally considered as a balanced measure that can be used even though the classes are of very different sizes.

Likewise, F1-score is the harmonic average of the precision and recall and is generally considered another balanced measure when the dataset is imbalanced. Since F1-score considers harmonic average, it is considered to provide an appropriate score to the model rather than an arithmetic mean. From Table 7, it is clear that based on MCC and F1-score AIRBP outperforms RBPPred by 9.53% and 6.75%.

#### Performance Comparison with Existing Approaches on the Independent Test Set

##### Performance Comparison with RBPPRed

In this section, we further compare the performance of AIRBP with RBPPred predictor on three different independent test sets, Human, S. cerevisiae and A. thaliana. Here, we only report the comparison of AIRBP with RBPPred because RBPPred is the top-performing sequence-based predictor of RBPs in the literature. As reported, RBPPred provides much better performance than SPOT-seq (Zhao, et al., 2011) and RNApred (Kumar, et al., 2011) predictors, which are the only two additional sequence-based methods that can be accessed either through a web server or code is publicly available for download. To perform independent testing, we first train AIRBP on a complete benchmark dataset and subsequently test it on three different independent test sets, Human, S. cerevisiae and A. thaliana. The predictive results of AIRBP and RBPPred on three different independent test sets are compared in Table 8. Table 8 indicates that AIRBP achieves an improvement of 9.32% in SN, 4.54% in ACC, 4.19% in F1-score and 8.50% in MCC over RBPPred on Human test set. Likewise, AIRBP achieves an improvement of 9.51% in SN, 4.41% in ACC, 3.52% in F1-score and 8.23% in MCC over RBPPred on S. cerevisiae test set. Furthermore, while testing on A. thaliana, AIRBP achieves an improvement of 6.61% in SN, 5.34% in ACC, 4.28% in PR, 3.03% in F1-score and 10.61% in MCC over RBPPred approach.

**Table 8.**
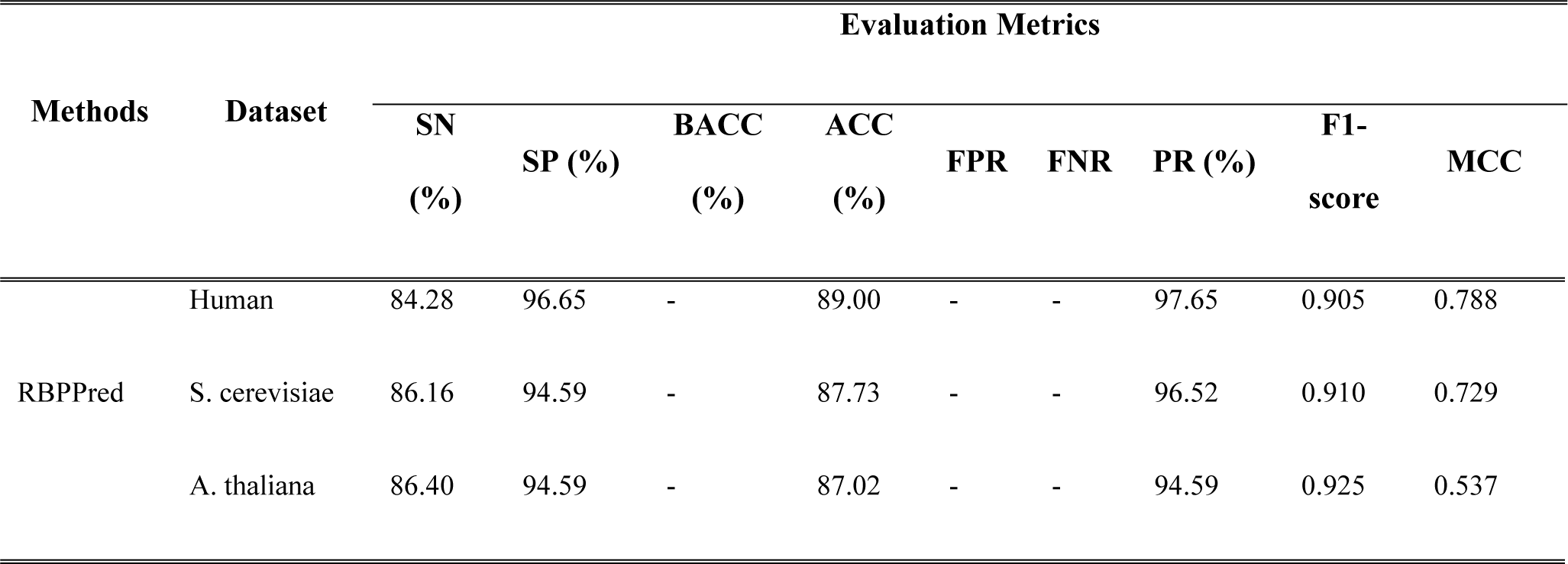
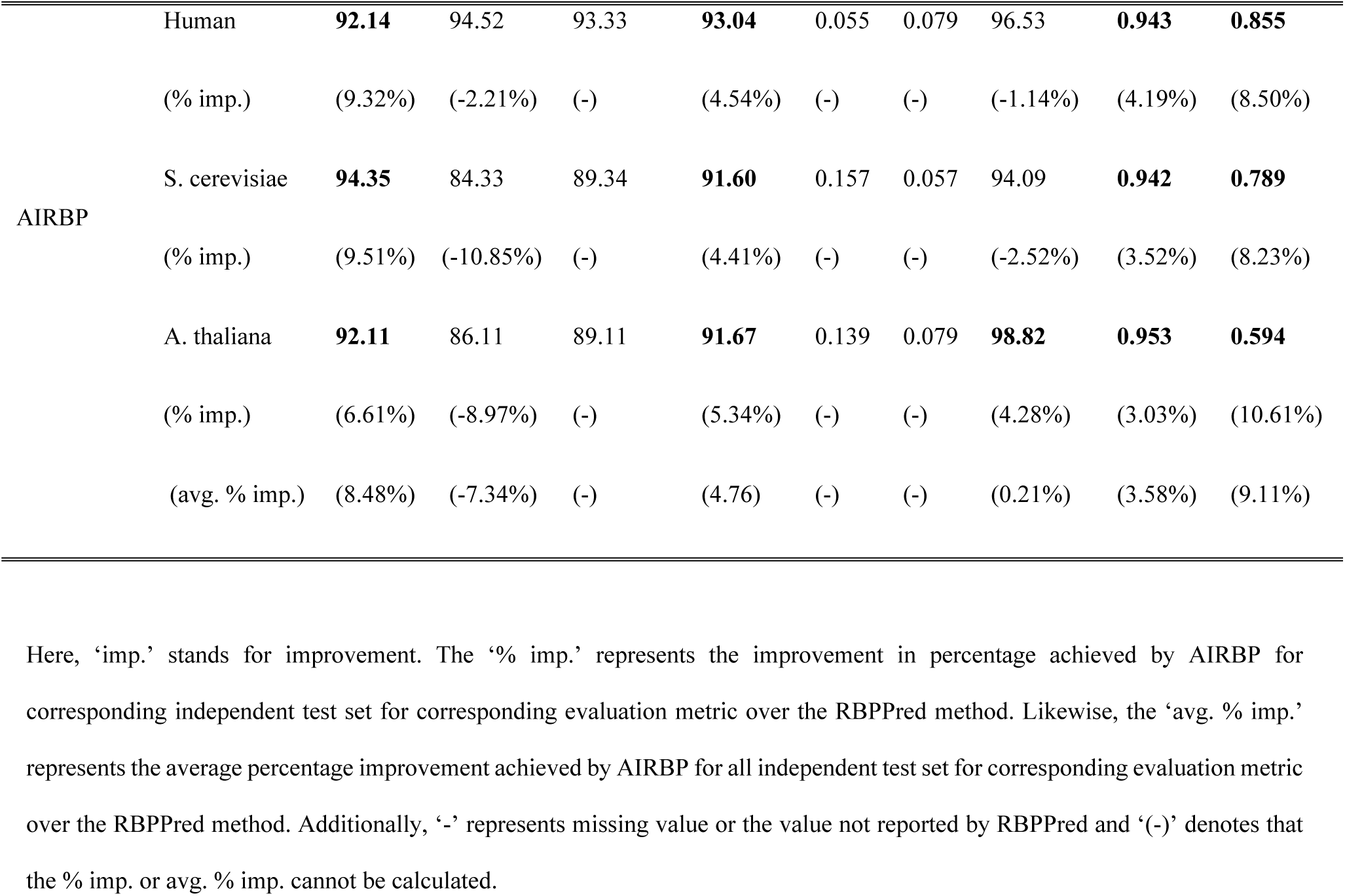
Comparison of AIRBP with the existing method on independent test sets.

Moreover, while analyzing the average percentage improvement over all the independent test sets AIRBP attains improvement of 8.48% in SN, 4.76% in ACC, 0.21% in PR, 3.58% in F1-score and 9.11% in MCC over RBPPred. Besides, RBPPred seems to be 7.34% better in an average over three test sets in terms of SP (i.e. predicting negative samples or non-RBPs) over AIRBP. However, AIRBP provides a 0.21% improvement in an average over three test sets in terms of PR over RBPPred.

Additionally, as stated above, for the imbalanced dataset the F1-score and MCC are widely used as a balanced measure between sensitivity and specificity. Our predictor, AIRBP shows consistent improvement in F1-score and MCC over RBPPred for all three independent test sets. Specifically, AIRBP provides 4.19% and 8.05% improvement in F1-score and MCC, respectively over RBPPred while tested on the Human test set. Similarly, AIRBP shows 3.52% and 8.23% improvement in F1-score and MCC, respectively, over RBPPred on S. cerevisiae as well as 3.03% and 10.61% improvement in F1-score and MCC, respectively over RBPPred on A. thaliana test set. Finally, based on an average percentage improvement (calculated over three different datasets) in F1-score and MCC, AIRBP outperforms RBPPred by 3.58% and 9.11%.

#### Performance Comparison with Deep-RBPPred and TriPepSVM

In this section, we compare the performance of AIRBP with two additional predictors, Deep-RBPPred (Zheng, et al., 2018) and TriPepSVM (Bressin, et al., 2019) on three different independent test sets, Human, S. cerevisiae and A. thaliana. To compare, first, we downloaded both the software Deep-RBPPred and TriPepSVM that are openly available from the internet. Next, we ran both Deep-RBPPred and TriPepSVM on the three independent test datasets, ATH, SC, and Human, respectively. The details of the process adopted for the comparison and the results obtained are provided below:

#### Comparison with Deep-RBPPred

In Deep-RBPPred software, to make predictions, users can either choose a model trained with the balanced dataset (balanced model) or a model trained with the imbalanced dataset (imbalanced model). In our implementation, we ran both balanced and imbalanced model of Deep-RBPPred on the three independent test datasets. Table 9 shows the comparison between the proposed method, AIRBP and an existing method Deep-RBPPRed on three independent test datasets. From Table 9, we can see that AIRBP consistently results in a higher number of True Positive (TP) count, which indicates that AIRBP is better than Deep-RBPPred in identifying RNA-binding proteins correctly, which is the primary objective of this work. The number of RNA-binding proteins correctly predicted by AIRBP is 15 counts less than the Deep-RBPPred Balanced model for the Human test dataset. However, the number of RNA-binding proteins correctly predicted by AIRBP is 64 counts higher than Deep-RBPPred Imbalanced for the same Human test dataset. Moreover, AIRBP accurately predicts 8 and 48 count of additional RNA-binding proteins compared to DeepRBPPred Balanced and Deep-RBPPred Imbalanced model for the ATH dataset, respectively. Similarly, AIRBP correctly predicts 9 and 39 count of other RNA-binding proteins compared to DeepRBPPred Balanced and Deep-RBPPred Imbalanced model for the SC dataset, respectively.

**Table 9.**
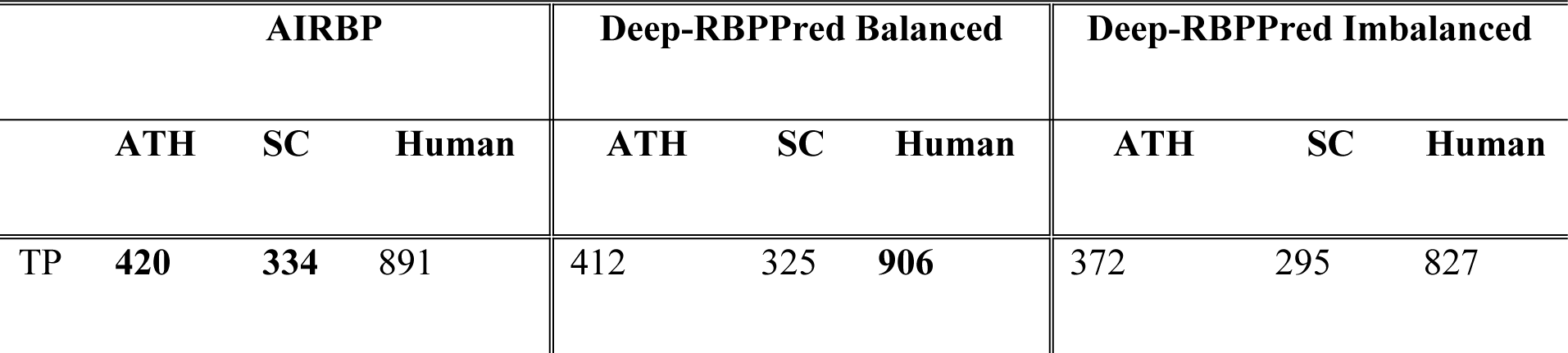
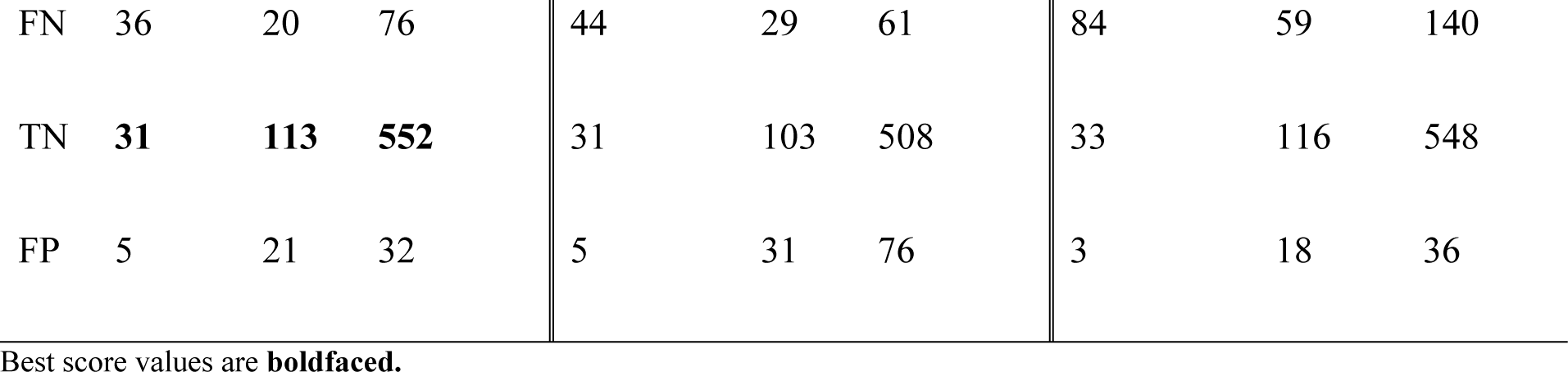
Comparison between AIRBP and Deep-RBPPred on three independent test datasets.

| AIRBP |  |  |  | Deep-RBPPred Balanced |  |  | Deep-RBPPred Imbalanced |  |  |
| --- | --- | --- | --- | --- | --- | --- | --- | --- | --- |
|  | ATH | SC | Human | ATH | SC | Human | ATH | SC | Human |
| TP | 420 | 334 | 891 | 412 | 325 | 906 | 372 | 295 | 827 |
| FN | 36 | 20 | 76 | 44 | 29 | 61 | 84 | 59 | 140 |
| TN | **31** | **113** | **552** | 31 | 103 | 508 | 33 | 116 | 548 |
| FP | 5 | 21 | 32 | 5 | 31 | 76 | 3 | 18 | 36 |

Best score values are **boldfaced**.

Moreover, we compare the performance of AIRBP with Deep-RBPPred on the three new independent test datasets (ATH-Filtered, SC-Filtered, and Human-Filtered) that we created by filtering sequences through similarity search using CD-Hit program in Table 10. We created ATH-Filtered, SC-Filtered, and Human-Filtered by filtering similar sequences that are more than 25% similar between training and ATH, SC, and Human datasets, respectively using CD-Hit program. Table 10 shows that AIRBP attains the highest count of true positive (TP) or in other words, predicts the highest number of RNA-binding proteins correctly compared to Deep-RBPPred Balanced, Deep-RBPPred Unbalanced and RBPPred for both SC and Human datasets. Further, we can see that for the ATH dataset the TP count of AIRBP is just 1 lower than Deep-RBPPred Balance, whereas 1 and 4 counts are higher than Deep-RBPPred Unbalanced and RBPPred, respectively. Again, we are focusing on TP counts because our major goal here is to accurately predict RNA-binding proteins. In summary, the results presented in Tables 8 and 9 show that AIRBP is a better predictor compared to both RBPPred and Deep-RBPPred in the majority of the cases.

**Table 10.**
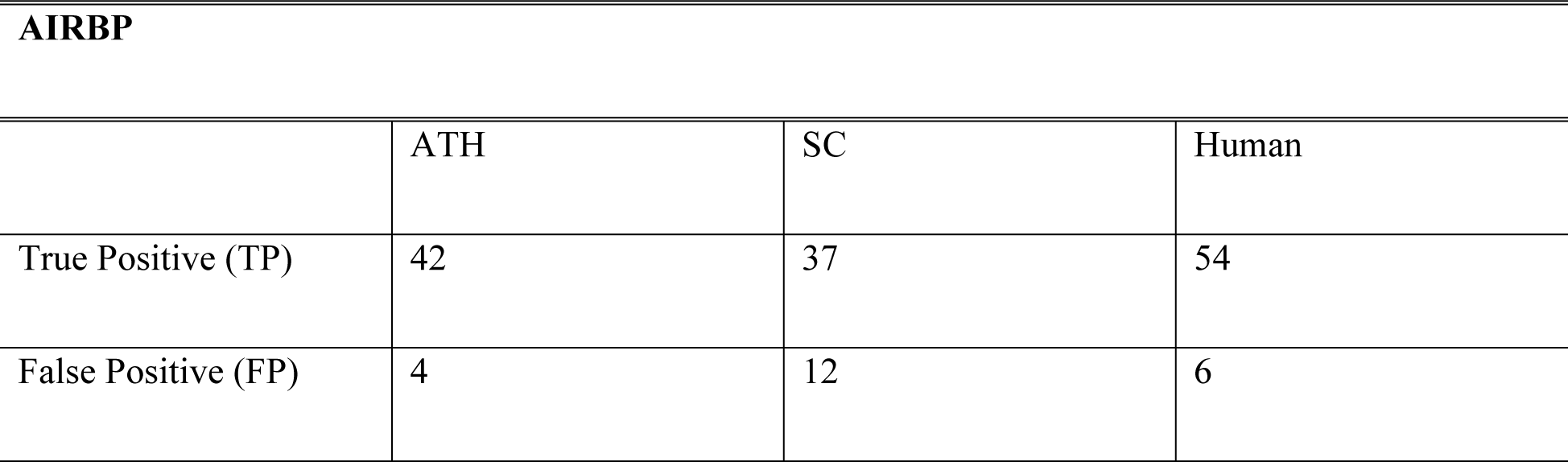
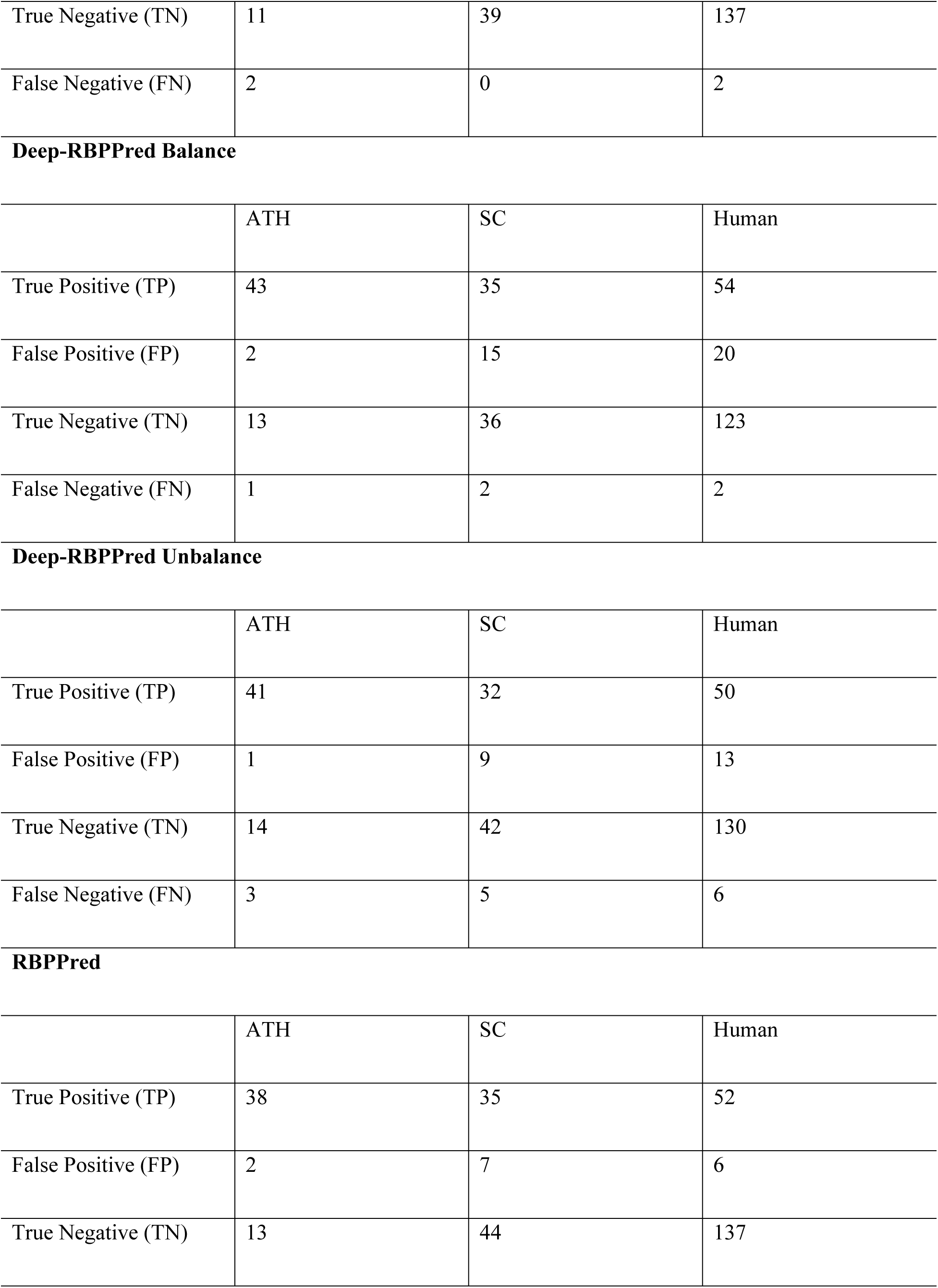
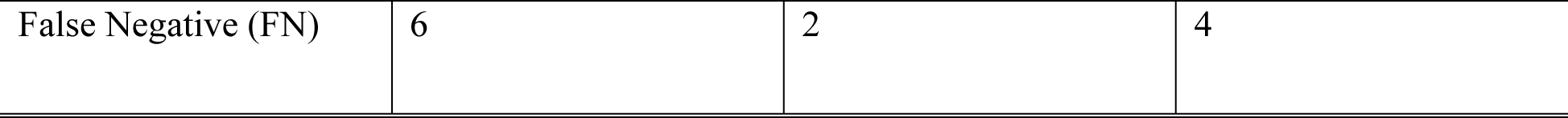
Comparison of AIRBP with RBPPred and Deep-RBPPred on the three independent test datasets ATH-Filtered, SC-Filtered, and Human-Filtered.

| **AIRBP** |  |  |  |
| --- | --- | --- | --- |
|  | ATH | SC | Human |
| True Positive (TP) | 42 | 37 | 54 |
| False Positive (FP) | 4 | 12 | 6 |
| True Negative (TN) | 11 | 39 | 137 |
| False Negative (FN) | 2 | 0 | 2 |
| **Deep-RBPPred Balance** |  |  |  |
|  | ATH | SC | Human |
| True Positive (TP) | 43 | 35 | 54 |
| False Positive (FP) | 2 | 15 | 20 |
| True Negative (TN) | 13 | 36 | 123 |
| False Negative (FN) | 1 | 2 | 2 |
| **Deep-RBPPred Unbalance** |  |  |  |
|  | ATH | SC | Human |
| True Positive (TP) | 41 | 32 | 50 |
| False Positive (FP) | 1 | 9 | 13 |
| True Negative (TN) | 14 | 42 | 130 |
| False Negative (FN) | 3 | 5 | 6 |
| **RBPPred** |  |  |  |
|  | ATH | SC | Human |
| True Positive (TP) | 38 | 35 | 52 |
| False Positive (FP) | 2 | 7 | 6 |
| True Negative (TN) | 13 | 44 | 137 |
| False Negative (FN) | 6 | 2 | 4 |

#### Comparison with TriPepSVM

Likewise, to compare AIRBP with TriPepSVM, we ran TriPepSVM on three independent test datasets. While running TriPepSVM, we discovered that it requires Uniprot taxon id, which is by default set to 9606 (for humans). This indicates that TriPepSVM must have been trained based on the species-wise dataset. For a quick check, we ran TriPepSVM on the ATH dataset with Uniprot taxon id, ATH, and Human, respectively, and with no surprise, we found that TriPepSVM resulted in inferior performance, while we ran it on ATH dataset with Human taxon id. So, one of the limitations of TriPepSVM is that it does not apply to the datasets of new species. The performance of TriPepSVM while using both ATH and Human taxon id is shown in Table 11. From Table 11, we can conclude that TriPepSVM is not a generic method that can be applied to any species. Instead, it is strongly dependent on the Uniprot taxon id as well as it will only perform well for particular species but not for any species. Therefore, we would like to highlight that the comparison between AIRBP and TriPepSVM is not an apple to apple comparison.

**Table 11.**
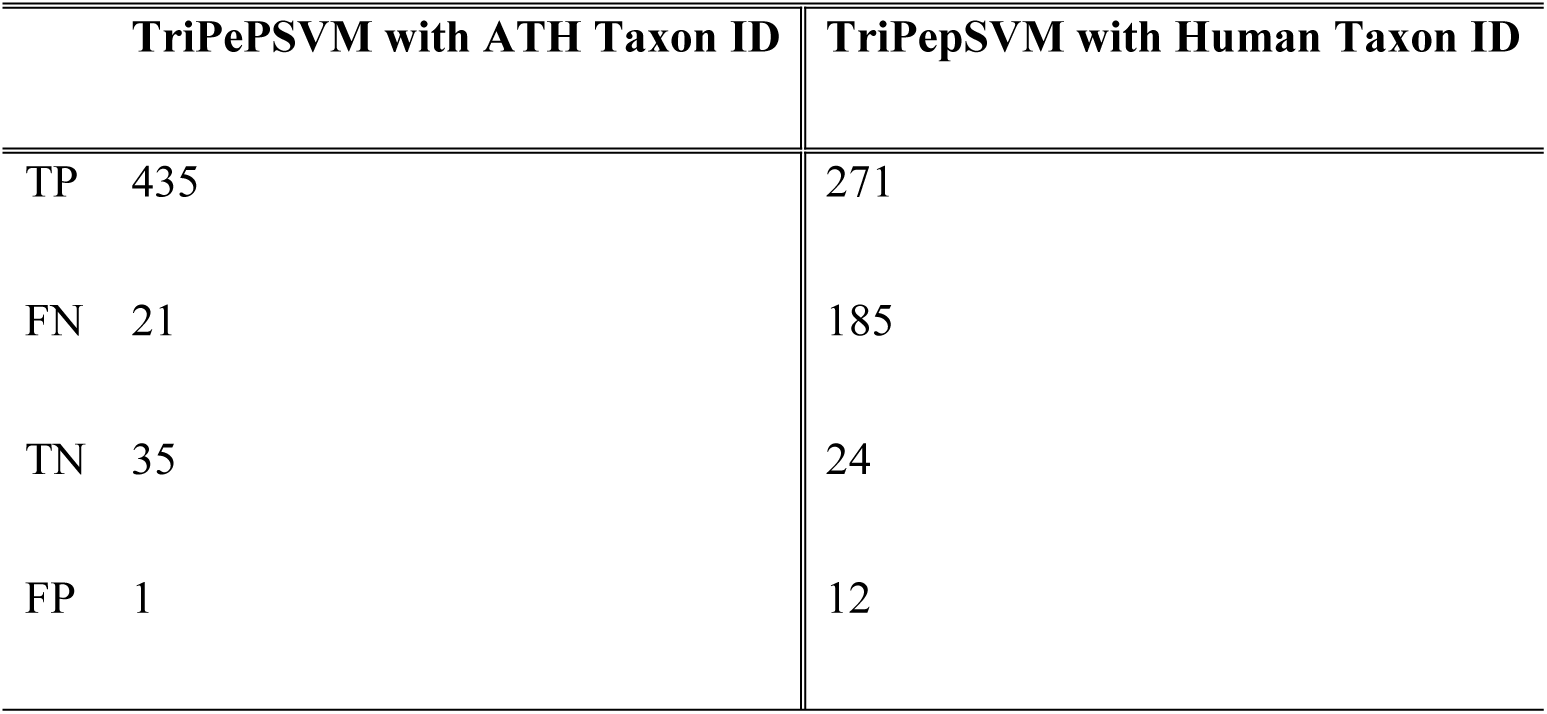
Performance of TriPepSVM on ATH dataset while using ATH and Human taxon id, respectively.

Further, Table 12 shows the comparison between the proposed method, AIRBP and an existing method TriPepSVM on three independent test datasets.

**Table 12.** Comparison between AIRBP and TriPepSVM on three independent test datasets.

| AIRBP |  |  |  | TriPepSVM |  |  |
| --- | --- | --- | --- | --- | --- | --- |
|  | ATH | SC | Human | ATH | SC | Human |
| TP | 420 | 334 | 891 | 435 | 42 | 953 |
| FN | 36 | 20 | 76 | 21 | 312 | 14 |
| TN | 31 | 113 | 552 | 35 | 135 | 462 |
| FP | 5 | 21 | 32 | 1 | 0 | 122 |

From Table 12, it is evident that even though TriPepSVM performs better than AIRBP on two independent test datasets, ATH and Human, the performance of TriPepSVM is exceptionally poor on SC test set. Again, better performance of TriPepSVM on some test set whereas, exceptionally poor performance on another test set also indicates that TriPepSVM is not a generic tool rather, it is trained on dataset of specific species and will only perform well for that particular species. On the contrary, AIRBP shows a very consistent performance on all the test datasets and therefore, AIRBP is a generic tool that can be applied for the prediction of RNA-binding proteins that may belong to any type of species. In other words, AIRBP is not tied to any particular species and can identify RNA-binding proteins from any species.

Further, the poor performance of TriPepSVM on SC test dataset encouraged us to perform additional analysis to identify if combining TriPepSVM with AIRBP would correct issues with TriPepSVM. Our first analysis involved combining TriPepSVM within the stacking framework of AIRBP. We added TriPepSVM as one of the base-layer classifiers into our stacking framework. Specifically, we ran TriPepSVM on our training dataset to collect prediction probabilities and added these probabilities as a feature vector to re-train the meta-layer classifier, XGBoost of our stacking framework. In Table 13, we compare the performance of AIRBP with the new stacking framework created by adding TriPepSVM at the base layer of AIRBP, we represent this new stacking framework as AIRBP+TriPepSVM.

**Table 13.** Comparison of AIRBP with AIRBP+TriPepSVM on the benchmark dataset through a 10-fold CV.

| **Metric/Methods** | **TriPepSVM+AIRBP** | **AIRBP** |
| --- | --- | --- |
| SN (%) | 90.57 | 90.17 |
| SP (%) | 97.57 | 97.44 |
| BACC (%) | 94.06 | 93.80 |
| ACC (%) | 95.58 | 95.38 |
| FPR | 0.024 | 0.026 |
| FNR | 0.094 | 0.098 |
| PR (%) | 93.65 | 93.31 |
| F1-score | 0.921 | 0.917 |
| MCC | 0.890 | 0.885 |

From Table 13, we can see that adding TriPepSVM as the base-layer classifier resulted in a slight improvement in the performance of all the performance measures used in our study. Particularly, ACC, F1-score and MCC of AIRBP+TriPepSVM framework increased from 95.38, 0.917, and 0.885 to 95.58, 0.921, and 0.890, respectively.

Next, we tested the performance of AIRBP+TriPepSVM on three independent test datasets. In Table 14, we compare the performance of AIRBP with the AIRBP+TriPepSVM framework on three independent test datasets. From Table 14, we can see that the performance of the AIRBP+TriPepSVM framework is slighter better than AIRBP for the Human test set. However, AIRBP outperforms AIRBP+TriPepSVM on SC and ATH test sets. Notably, for the Human test set, AIRBP+TriPepSVM results in a thinly better MCC of 0.858 compared to MCC of 0.855 by AIRBP. However, for SC and ATH test set, AIRBP results in significantly better with MCC of 0.789 and 0.594 compared to MCC of 0.465 and 0.54 by AIRBP+TriPepSVM. Overall, the comparison of AIRBP with TriPepSVM and AIRBP+TriPepSVM indicates that TriPepSVM suffers from an inconsistency issue as it works better on one dataset, whereas it performs very poorly on another dataset. This inconsistency can be overcome by adding TriPepSVM as a base layer in AIRBP to produce the AIRBP+TriPepSVM framework. However, the performance of AIRBP+TriPepSVM on independent test datasets is still lower compared to AIRBP. This helps us conclude that AIRBP performs better compared to Deep-RBPPred in the majority of the cases. Moreover, we would also like to highlight that comparison between AIRBP and TriPepSVM is not an apple-to-apple comparison as TriPepSVM is trained based on specific species and is not a generic method as AIRBP.

**Table 14.** Comparison of AIRBP, TriPepSVM, and AIRBP+TriPepSVM on three independent test datasets.

| Methods | Dataset | Evaluation Metrics |  |  |  |  |  |  |  |  |
| --- | --- | --- | --- | --- | --- | --- | --- | --- | --- | --- |
|  |  | SN | SP | BACC | ACC | FPR | FNR | PR | F1-score | MCC |
|  |  | (%) | (%) | (%) | (%) |  |  | (%) |  |  |
| AIRBP+<br>TriPepSVM | Human | 92.87 | 93.84 | 93.35 | 93.23 | 0.062 | 0.071 | 96.14 | 0.945 | 0.858 |
|  | SC | 58.48 | 93.28 | 75.88 | 75.88 | 0.067 | 0.415 | 95.83 | 0.726 | 0.465 |
|  | ATH | 87.50 | 94.44 | 90.97 | 88.08 | 0.056 | 0.125 | 99.50 | 0.931 | 0.54 |
| AIRBP | Human | 92.14 | 94.52 | 93.33 | 93.04 | 0.055 | 0.079 | 96.53 | 0.943 | 0.855 |
|  | SC | 94.35 | 84.33 | 89.34 | 91.60 | 0.157 | 0.057 | 94.09 | 0.942 | 0.789 |
|  | ATH | 92.11 | 86.11 | 89.11 | 91.67 | 0.139 | 0.079 | 98.82 | 0.953 | 0.594 |

The above comparison of results indicates that the proposed method, AIRBP outperforms the existing methods and is a very promising predictor. We believe that this comprehensive investigation of the stacking based machine learning framework and features in predicting RNA binding proteins might be useful for future proteomics studies.

## Conclusions

In this work, we constructed a stacking based machine learning framework, called AIRBP, for the prediction of RNA-binding proteins directly from the protein sequence. To improve the prediction accuracy of RNA-binding proteins, we have investigated and used various feature extraction and encoding techniques, different feature selection techniques along with an advanced machine learning technique called stacking. We extracted multiple features, including evolutionary information, physiochemical properties, and disordered properties and applied different encoding techniques such as composition, transition and distribution, conjoint triad, PSSM distance transformation, and residue-wise contact energy matrix transformation to encode the protein sequence in terms of features. Next, we applied two different feature selection techniques, incremental feature selection and evolutionary algorithm based feature selection to identify the relevant features as well as to reduce the feature space significantly. Next, only the relevant features are used to train the ensemble of predictors at the first-level (i.e. base-layer) of the stacking framework. Then, the prediction probabilities from the first-level predictors are combined with the originally selected features and used to train the predictor at the second-level (i.e. meta-layer) of the stacking framework. Finally, the AIRBP stacking framework achieves a 10-fold CV accuracy, F1-score, and MCC of 95.38%, 0.917 and 0.885 respectively, on the benchmark dataset. While performing the independent test, AIRBP achieves an accuracy, F1-score, and MCC of 93.04%, 0.943 and 0.855, for Human test set; 91.60%, 0.942 and 0.789 for S. cerevisiae test set; and 91.67%, 0.953 and 0.594 for A. thaliana test set, respectively. These promising results indicate that the stacking framework helps improve the accuracy significantly by reducing the generalization error. Furthermore, in comparison with the top-performing method, RBPPred, AIRBP achieves 3.26%, 6.75% and 9.53% improvement in terms of accuracy, F1-score and MCC respectively, based on a benchmark dataset. F1-score and MCC are two widely used measures for the imbalanced dataset. Moreover, the average percentage improvement, calculated over three different independent test sets, AIRBP outperforms RBPPred by 4.76%, 3.58% and 9.11% in terms of accuracy, F1-score, and MCC, respectively. These outcomes help us summarize that the AIRBP can be effectively used for accurate and fast identification and annotation of RNA-binding proteins directly from the protein sequence and can provide valuable insights for treating critical diseases.

## Acknowledgments

We would like to thank the anonymous reviewers.

## Author Contributions

Conceived and designed the experiments: AM, RK MTH. Performed the experiments: AM, RK. Analyzed the data: AM, RK. Contributed reagents/materials/analysis tools: MTH. Wrote the paper: AM, RK MTH.

